# Towards an individualised neural assessment of receptive language in children

**DOI:** 10.1101/566752

**Authors:** Selene Petit, Nicholas A. Badcock, Tijl Grootswagers, Anina N. Rich, Jon Brock, Lyndsey Nickels, Denise Moerel, Nadene Dermody, Shu Yau, Elaine Schmidt, Alexandra Woolgar

## Abstract

**Purpose:** We aimed to develop a non-invasive neural test of language comprehension to use with non-speaking children for whom standard behavioural testing is unreliable (e.g., minimally-verbal autism). Our aims were three-fold. First, we sought to establish the sensitivity of two auditory paradigms to elicit neural responses in individual neurotypical children. Second, we aimed to validate the use of a portable and accessible electroencephalography (EEG) system, by comparing its recordings to those of a research-grade system. Third, in light of substantial inter-individual variability in individuals’ neural responses, we assessed whether multivariate decoding methods could improve sensitivity.

**Methods:** We tested the sensitivity of two child-friendly covert N400 paradigms. Thirty-one typically developing children listened to identical spoken words that were either strongly predicted by the preceding context or violated lexical-semantic expectations. Context was given by a cue word (Experiment 1) or sentence frame (Experiment 2) and participants either made an overall judgement on word relatedness or counted lexical-semantic violations. We measured EEG concurrently from a research-grade system, Neuroscan’s SynAmps2, and an adapted gaming system, Emotiv’s EPOC+.

**Results:** We found substantial inter-individual variability in the timing and topology of N400-like effects. For both paradigms and EEG systems, traditional N400 effects at the expected sensors and time points were statistically significant in around 50% of individuals. Using multivariate analyses, detection rate increased to 88% of individuals for the research-grade system in the sentences paradigm, illustrating the robustness of this method in the face of inter-individual variations in topography.

**Conclusions:** There was large inter-individual variability in neural responses, suggesting inter-individual variation in either the cognitive response to lexical-semantic violations, and/or the neural substrate of that response. Around half of our neurotypical participants showed the expected N400 effect at the expected location and time point. A low-cost, accessible EEG system provided comparable data for univariate analysis but was not well suited to multivariate decoding. However, multivariate analyses with a research-grade EEG system increased our detection rate to 88% of individuals. This approach provides a strong foundation to establish a neural index of language comprehension in children with limited communication.

## 1 Introduction

Language is a crucial part of everyday life and something we often take for granted. In cases where people cannot speak or reliably communicate, it can be difficult to assess whether an individual is able to understand spoken language. Examples include disorders of consciousness, minimally-verbal autism, or cerebral palsy (Giacino & Smart, 2007; Harrison & Connolly, 2013; Tager-Flusberg & Kasari, 2013). In these cases, individuals may score poorly on standardised or behavioural tests of language despite intact language processing. For example, recent research on minimally-verbal autistic children reports intact neural markers of lexico-semantic processing despite an absence of spoken language (Cantiani et al., 2016; DiStefano et al., 2019). This highlights the need for alternatives to standard measures of receptive language. Such a method could transform both individual assessment and treatment, and our scientific understanding of the cognitive profile of these individuals.

Previous research has used indirect measures to investigate individual differences in language processing in neurotypical and clinical populations. For example, eye-tracking has been used to study inter-individual differences in word processing (Farris-Trimble & McMurray, 2013). One commonly used eye-tracking paradigm, preferential-looking tasks, consists of presenting participants with a spoken word, and two visual displays, one of them matching the auditory description, and computing the time they spend looking at each visual display. Looking-time to the matching display reveals processing of the auditory word (Fernald et al., 2008; Swingley, 2011). This logic has been used to assess language processing in minimally-verbal populations such as young infants (Brandone et al., 2007; Seidl et al., 2003) and children with motor impairments (Cauley et al., 1989). However, eye tracking techniques are not suitable in populations where gaze fixation or eye-movements may be impaired, such as in autism (Riby & Doherty, 2009; Schmitt et al., 2014).

Non-invasive neuroimaging, such as electroencephalography (EEG), may offer an opportunity to measure language understanding in the absence of reliable behavioural or eye-movement responses. EEG is a passive way to record neural responses, and offers the opportunity to measure language understanding in the absence of reliable behavioural responses. In particular, the N400 event-related potential (ERP) is elicited by hearing or reading words, with the N400 being larger when the words violate the semantic or predictive context in which they are presented (e.g., Kutas, 1993; Kutas & Hillyard, 1980; Kutas & Federmeier, 2011). For instance, the word “door” elicits a larger N400 ERP when it is presented in the sentence “The clouds are high up in the *door*”, compared to the sentence “I had no key to open the *door*”. This difference in the N400 amplitude is called the N400 effect. It is well-documented and has been recorded in groups of adults, children, and special populations (for a comprehensive review see Kutas & Federmeier, 2011), making it a strong candidate for assessing lexical-semantic processing. Accordingly, the N400 has previously been used as a neural index of linguistic processing in special populations, such as cerebral palsy (Byrne et al., 1999, 1995), traumatic brain injury (Connolly et al., 1999), stroke (D’Arcy et al., 2003), autism (Cantiani et al., 2016; Coderre et al., 2019; DiStefano et al., 2019), and disorders of consciousness (Schoenle & Witzke, 2004; Steppacher et al., 2013).

However, even in neurotypical populations, few studies have systematically assessed the ability to reliably detect the N400 effect in individuals (Beukema et al., 2016; Cruse et al., 2014; Rohaut et al., 2015), and to our knowledge, only one has reported them for individual children (Cantiani et al., 2016; albeit with only ten participants). To date, the sparse results on neurotypical individuals suggest that, despite robust effects at the group level, current N400 paradigms allow to detect N400 effect to semantic manipulations in around half of the neurotypical individuals. This has important implications both for neuro-cognitive understanding of lexico-semantic processing, and for the possibility of using N400 as a future neural marker of language processing in non-verbal individuals. Because of the sparsity of results in individual children and the variable N400 effect detection rate in previous adult studies, our first aim was to add to the knowledge base by establishing the sensitivity of two auditory paradigms for detecting lexico-semantic processing in individual neurotypical children.

In our first experiment, neurotypical children listened to pairs of spoken words which were either normatively associated (e.g., “arm – *leg*”) or unrelated (e.g., “boat – *leg*”). In our second experiment, children listened to spoken sentences with either a congruent completion (i.e., “she wore a necklace around her *neck*”) or an incongruent/anomalous completion (i.e., “the princess may someday become a *neck*”). These paradigms elicit strong N400 effects in groups of adults and children (Borovsky et al., 2012; Friedrich & Friederici, 2005; Kiang et al., 2013; Rämä et al., 2013; Torkildsen et al., 2007), so we predicted that the ERP evoked by the identical spoken word token (in our example, *neck*) would vary according to semantic context, and asked whether we could detect this difference in individual children. To make the paradigms suitable for children we created game-like tasks where children encountered friendly or evil aliens.

Our second aim, with a view to future clinical applications, was to validate an accessible EEG system that avoids many of the typical setup inconveniences. A traditional 32-channel, gel-based EEG system takes around 35 minutes to setup and is somewhat uncomfortable, involving rubbing the participant’s scalp with gel in order to bridge the EEG electrodes to the skin. In addition, most typical EEG setups are not portable, with extensive wiring compelling participants to remain seated, and signals are best recorded in an electrically shielded room, which can be intimidating for some subjects. Although previous research has successfully used laboratory EEG systems to test special populations such as autistic individuals (Coderre et al., 2019; McCleery et al., 2010) and individuals with Rett syndrome (Laan & Vein, 2002), alternative portable solutions may allow the inclusion of more individuals in research. For example, the laboratory environment may be too intimidating for some autistic children, with portable solutions allowing testing in a more familiar environment. Recently, more accessible and more portable EEG systems have become available, one of which has been adapted and validated for the measurement of auditory ERPs in adults (Badcock et al., 2013; de Lissa et al., 2015a), children (Badcock et al., 2015), and autistic children (Yau et al., 2015). The Emotiv EPOC+ system, hereafter referred to as EPOC+, was originally designed for gaming purposes, and consists of a wireless headset with 14 electrodes that connect to the scalp via saline solution-soaked cotton-rolls. The setup is fast (approx. 5-10 minutes) and it is not necessary to rub the scalp. This system is also low in cost compared with research-grade systems and is wireless and portable, allowing its use outside of the laboratory (e.g., in homes or schools).

Although the EPOC system (the predecessor of EPOC+) has been validated against research-grade systems for recording early ERPs, such as auditory ERPs (Badcock et al., 2015, 2013; Barham et al., 2017), and face-sensitive N170 (de Lissa et al., 2015b), studies on later components such as the P300 have yielded less consistent results. Vos et al. (2014) report similar performance of an Emotiv amplifier compared to a research-grade amplifier, and Elsawy et al. (2014) found acceptable results when using a classifier on P300 EPOC data, but Duvinage et al. (2013) report that the EPOC recorded a significantly noisier signal compared to the research-grade ANT system (Advanced Neuro Technology, ANT, Enschede, The Netherlands). To our knowledge, no studies have tested it on N400 ERPs. Here, we tested the fidelity of the adapted Emotiv EPOC+ EEG system against data recorded concurrently from a research-grade Neuroscan system during the experimental tasks.

Our third aim was to assess whether we could improve detection of lexico-semantic processing at the individual level by using more sensitive analytical methods. Traditional univariate analyses of ERPs usually require an *a priori* choice of electrodes and time points of interest. However, when testing individual participants, especially children and special populations, this *a priori* knowledge may not be available. In contrast, multivariate pattern analyses (MVPA) allow for consideration of multiple electrodes at once, removing the requirement for *a priori* knowledge of topology without introducing multiple comparisons. Accordingly, we compared our detection rate of ERP differences between semantic conditions using both typical univariate N400 analyses and MVPA. For MVPA, we trained a linear classifier to discriminate between the two semantic conditions (congruent and incongruent) on individual-subject EEG data. This targets the information contained in the pattern of activation across sensors, making it robust to individual differences in signal direction and topology (Grootswagers et al., 2017; Haynes, 2015; Hebart & Baker, 2018).

To pre-empt our results, we found both robust group-level univariate N400 effects and substantial inter-subject variability in topology and time course of response. The individual subject detection rate was around 50% in both paradigms, for both research-grade and EPOC+ systems, using classical analysis. In the sentence paradigm, MVPA increased the detection rate to 88% but only for the research-grade EEG system. The data suggest heterogeneity in the neural responses to lexico-semantic processing in neurotypical children and may help direct future development of paradigms for assessing language comprehension in non-speaking children.

## 2 Experiment 1: associated word pairs

### 2.1 Methods

#### 2.1.1 Participants

Sixteen children were recruited using the Neuronauts database of the Australian Research Council Centre of Excellence in Cognition and its Disorders. All participants were native English speakers and had non-verbal reasoning and verbal abilities within the normal range as measured by the matrices section of the Kaufman Brief Intelligence Test, Second Edition (K-BIT 2, Kaufman & Kaufman, 2004) and the Peabody Picture Vocabulary Test— 4th Edition (PPVT – 4, Dunn & Dunn, 2007). Participants received AUD25 for their participation, as well as a sticker and a certificate. The data from one participant were excluded due to technical issues during recording. The final set of data thus came from 15 participants (age range: 6 to 12 years old, *M=9*.*2, SD=2*.*6*, 4 male and 11 female). This study was approved by the Macquarie University Human Research Ethics Committee (Reference number: 5201200658). Participants’ parents or guardians provided written consent and the children provided verbal consent.

#### 2.1.2 Stimuli

Stimuli comprised 63 pairs of normatively associated words. Following Cruse et al. (2014), we began with word pairs taken from the Nelson et al. (1998) free association norms database. These norms comprise a large number of cue-target pairings, developed by asking participants to produce the first meaningfully or strongly associated word that comes to mind when presented with a particular cue. We initially chose pairs from the normative database with a forward associative strength (cue to target) greater than 0.5, meaning that more than 50% of the participants in the Nelson et al. norm-development study produced this target word in response to the cue. We included only pairs where the target was a noun, and where the cue and target were one syllable in length. We also included only words that had an age of acquisition rating of 8-years or less (Kuperman et al., 2012), meaning that these words were typically known by children of 8-years and above. It is possible that a few of our words were unknown to the younger participants. However, when selecting our stimuli, we chose to maximise the number of words fitting our criteria, acknowledging this may have resulted in some unknown words for some participants. We excluded any pairs where either the cue or target had a homophone (according to the N-watch database; Davis, 2005) with an age of acquisition of less than or equal to 10 years, and where the cue or target was not applicable to the Australian context (e.g., FUEL-GAS).

To minimise repetition across the stimulus set, we allowed each target word to appear in a maximum of two word pairs. The cue words were only used once as cues, but they could also appear up to twice as targets. Thus, the maximum number of repetitions of particular words across the entire list of related items was three (i.e., once as a cue, and twice as a target – this was the case for 13 words). For word pairs with singular and plural forms (e.g., GIRL-BOY and GIRLS-BOYS), only the pair with the strongest association was included. The final set of 63 pairs had a mean forward associative strength of 0.676 (see supplementary Table S1 for the list of stimuli). These word pairs formed the *related* condition.

We created a list of 63 *unrelated* word pairs by recombining the cue and target words from the related condition. Constructing the unrelated list in this way ensured a fully balanced design in which the cues and targets in the related and unrelated lists were identical (and therefore matched for word frequency, familiarity, phoneme length, etc.). We ensured that target words in the unrelated condition did not start with the same sound, rhyme, or have any semantic or associative connections with the cue or related target. In addition, we respected the grammatical number structure of the related word pair when choosing an unrelated target. For example, in creating an unrelated combination for a plural-singular pair (e.g., SUDS-SOAP), another singular target word was chosen (e.g., SUDS-ART).

Stimuli were digitally recorded by a female native Australian-English speaker, and the best auditory tokens, where the voice had a natural intonation and was not raspy, were selected using Praat software (Boersma, 2001). We used the same target tokens in the related and unrelated conditions so that there were no auditory differences to drive a differential EEG response to the two conditions. For each target, the related and the unrelated cue words were recorded close together in time, and were chosen to have approximately the same length, intensity, and voice quality as the target (mean stimulus length: 834 ms, SD: 121 ms, range: 590-1124 ms).

#### 2.1.3 EEG Equipment

We recorded simultaneously from two EEG systems in an electrically-shielded room. The research EEG system Neuroscan SynAmps2 (Scan version 4.3) Ag-AgCl electrodes were fitted to an elastic cap (Easy Cap, Herrsching, Germany) at 33 locations (Figure. 1), according to the international 10-20 system, including M1 (online reference), AFz (ground electrode), and M2. We measured vertical and horizontal eye movements with electrodes placed above and below the left eye and next to the outer canthus of each eye. Neuroscan was sampled at 1000Hz (down sampled to 500Hz during processing) with an online bandpass filter from 1 to 100Hz. We marked the onset of each sentence and target word using parallel port events generated using the Psychophysics Toolbox 3 extensions (Brainard, 1997; Kleiner et al., 2007; Pelli, 1997) in MATLAB.

**Figure 1.**
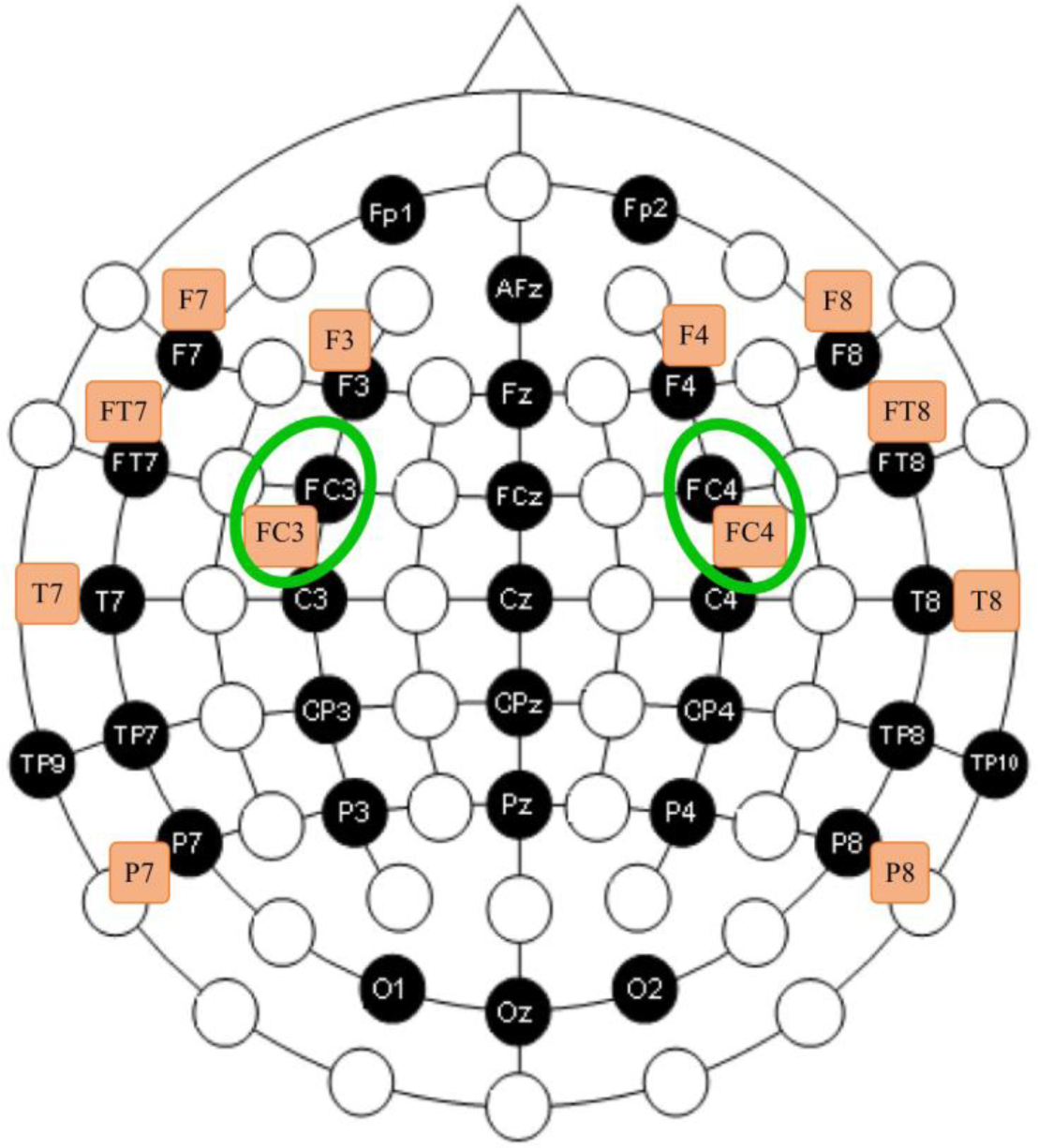
Electrode position on the scalp for the Neuroscan (black circles) and EPOC+ (orange rectangles) systems. The adjacent electrodes used to calculate correlations between the two systems are circled in green.

The EPOC+ is a wireless headset with flexible plastic arms holding 16 gold-plated sensors. In order to accommodate the concurrent setup with the Neuroscan system, we placed the EPOC+ sensors at the following scalp locations of the international 10-20 system (Klem et al., 1999; See Figure 1)^1^. M1 acted as the online reference, and M2 was a feed-forward reference that reduced external electrical interference. The signals from the other 14 channels were high-pass filtered online with a 0.16 Hz cut-off, pre-amplified and low-pass filtered at an 83 Hz cut-off. The analogue signals were then digitised at 2048 Hz. The digitised signal was filtered using a 5th-order sinc filter to notch out 50Hz and 60 Hz, low-pass filtered and down-sampled to 128 Hz (specifications taken from the EPOC+ system web forum). The effective bandwidth was 0.16–43 Hz.

To accurately time-lock the ERPs to the onset of the target word, we modified the EPOC+ system to incorporate event markers, following Badcock et al. (2015). We did this using a custom-made transmitter unit communicating with a custom receiver unit through infrared light (Thie, 2013). The transmitter box was connected to the audio output of the presentation computer. At the onset of each sentence and each target word, a tone of particular frequency (2400Hz for the sentence onset, 600Hz for a related target onset, and 1600Hz for an unrelated target onset) was sent to the transmitter unit through a separate audio channel. This in turn activated the receiver unit, which generated an electrical pulse in the O1 and O2 channels (from which we did not acquire neural data).

#### 2.1.4 Experimental Procedure

For each participant, we set up the Neuroscan system first and adjusted the impedances to under 5 kΩ. We then placed the EPOC+ system over the top, with cotton wool bridging scalp to sensor through custom slits in the EasyCap. EPOC+ impedances were adjusted to be below 20 kΩ in the TestBench software. Setup took up to 50 minutes, during which participants watched a DVD of their choice. Following setup, participants were seated in front of a 17-inch monitor, with speakers on both sides of the screen, at a viewing distance of about 1 metre. Before the main experiment, participants completed the PPVT 4, and after the EEG session, they completed the matrices section of the K-BIT 2.

Participants completed two EEG acquisition sessions of 20 minutes, separated by a 5-minute break. Each session included all 126 cue-target word pairs (63 related, 63 unrelated). The stimuli were presented using Psychophysics Toolbox 3 extensions (Brainard, 1997; Kleiner et al., 2007; Pelli, 1997) in MATLAB R2016B. The word pairs were presented in a pseudo-random order which was reversed in the second session. The order was optimised to minimise order bias in the sequence of related and unrelated trials, with the additional constraint that no more than 4 trials in a row were of either condition. The first session was divided into 17 blocks consisting of 5 to 15 trials, and the second section was divided into 11 blocks of 8 to 15 trials, as the participants were more familiar with the task during the second session.

To make the task more engaging for children, it was introduced in the context of a story. The child was told that they were listening to different aliens in an English-speaking competition. In the competition, aliens had to say pairs of related words (e.g., ‘boy-girl’, ‘left’-‘right’) and the children, and they had to rate each alien (1-5 stars) depending on how well it performed. We asked participants to listen carefully to each pair and to decide whether the two words were related or unrelated, then to give an overall judgment of the alien’s performance at the end of each block. Children were instructed to give between 1 star (if they produced mainly unrelated pairs) and 5 stars (if they said mainly related words). This encouraged the children to pay attention to, and make a covert decision about, the relatedness of the words in each pair. The choice of this covert task was motivated by the results from Cruse et al. (2014), who found that a covert design was more sensitive than a passive design where participants were asked to simply listen to the stimuli (58% detection rate for the covert task versus 0% for the passive task). According to the same study, an overt task would have been even more sensitive, but we anticipated that minimally-verbal individuals may not give reliable overt responses, whereas they may be able to perform the task covertly.

At the beginning of each block, an alien appeared on the screen, then moved behind a black “recording booth” in the middle of the screen. A light bulb was depicted on top of the box and lit up during each trial to encourage children to pay attention and to reduce eye movements during the trial. The light bulb lit up 500 ms before the cue word, remained lit until 1500 ms after the target word onset, then turned off. The interval between the cue and target was 1000 ms. After another 1500 ms, the next trial began. At the end of each block, the alien moved out of the box and participants were prompted to grade the alien on a five-point scale regarding its overall performance. At the end of the experiment, participants were shown the “winners” of the alien contest. The total time, including the EEG setup, was about 1hr 40 minutes.

#### 2.1.5 Offline EEG processing

We processed all EEG signals in EEGLAB (v13.4.4b, Delorme & Makeig, 2004) in MATLAB (R2014b). We first applied a band-pass two-way sinc FIR filter between 0.1 and 40Hz, allowing to have the same offline filtering procedure for both systems, then we cut the data into epochs from −100 ms to 1000 ms around target word onset. We ran an Independent Component Analysis (ICA) on all the epochs. Components with scalp distribution, frequency, and timing that corresponded to eye movements and eye blinks were removed from the Neuroscan data. In line with a previous EPOC+ study with children (Badcock et al., 2015), we could not identify any eye blink artefacts in the EPOC+ data. This may be because eye blinks were not consistent or strong enough to affect EPOC+ data. Alternatively, the ICA for the Neuroscan data could have benefited from the signal recorded by the Neuroscan electrodes recording eye movements. The EPOC+ did not have such electrodes so the ICA for the EPOC+ used only the scalp electrodes. The epochs were baseline corrected against the averaged signal from the 100 ms preceding the target onset. These data were then used for the decoding analyses. For univariate analyses, we further removed, from all electrodes, epochs with extreme values (±150μV) in any of the electrodes of interest (see below). As MVPA is more robust to noise in the data (Grootswagers et al., 2017b), and requires balanced data for valid statistical inference, we did not reject noisy trials for multivariate analyses. For the Neuroscan data, an average of 11 epochs (9%) for the related condition (SD = 7.75) and 10 epochs (8%) for the unrelated condition (SD = 7.56) were rejected. For the EPOC+ data, an average of 10 epochs (8%) for the related condition (SD = 7.72) and 9 epochs (7%) for the unrelated condition (SD = 6.11) were rejected.

#### 2.1.6 Group ERP analyses

The N400 is typically recorded over the centro-parietal regions of the brain. We therefore focused our univariate analyses on three electrodes sites of interest: Cz, as the N400 effect is reported to be the strongest in centro-parietal sites (Kutas & Federmeier, 2011), and FC3 and FC4, which are the closest channels to Cz that we can compare between the Neuroscan and EPOC+ systems.

We first verified that our paradigm evoked a classic N400 effect at a group level and tested whether this was detectable in both Neuroscan and EPOC+ systems. For trial-averaged waveforms, we ran group analyses with paired t-tests comparing the two conditions at each time point from 150 ms after stimulus onset for each of the five sensors of interest. We corrected for multiple-comparisons for each channel independently using a statistical temporal cluster extent threshold (Guthrie & Buchwald, 1991). Briefly, this method calculates the autocorrelation between consecutive time points of the ERP signal, for each channel. We can then determine the minimum number of consecutive time points that need to show a statistical difference in a one-tailed t-test between the related and unrelated conditions to be considered a significant cluster at p < .05 corrected (for details, see Guthrie & Buchwald, 1991). We used a one-tailed test because the direction of the N400 effect (a more negative response in the unrelated condition) was pre-specified. We restricted our statistical analyses to the time points after 150 ms to decrease the number of statistical tests computed, as the N400 is typically reported to occur later than 150 ms (Kutas & Federmeier, 2011).

We illustrated the topographic distribution of the N400 effect, based on the Neuroscan grand average data, by subtracting activation in the related condition from the unrelated condition and averaging over sequential 200 ms time windows, for ease of illustration (200 to 400 ms, 400 to 600 ms, 600 to 800 ms and 800 to 1000 ms).

#### 2.1.7 Single subject ERP analyses

Our next goal was to assess the sensitivity of our paradigm and EEG systems to detect N400 effects in individual children. For each individual and each system, we conducted first-level (single subject) analyses using independent samples t-tests between the two conditions at the electrodes Cz (for Neuroscan only), and FC3 and FC4 (for both systems), for each time point starting at +150 ms after the target onset (as we do not expect N400 effects to arise earlier than 150ms, i.e., Cruse et al., 2014). Although using independent t-tests is more conservative that paired t-tests across targets, we made this choice because the trial rejection performed during pre-processing left unbalanced (thus, not paired) trials. To correct for multiple comparisons, we again used the autocorrelation score to determine the temporal cluster extent threshold for each electrode in each participant independently (mean cluster extent threshold over participants and electrodes for Neuroscan: M = 132 ms, range = [70, 206], and for EPOC+: M = 58 ms, range = [47, 70]).

We illustrated the topographic distribution of the N400 effect in individuals based on the Neuroscan data by subtracting activation in the related from the unrelated condition and averaging over sequential 200 ms time windows, for ease of illustration (200 to 400 ms, 400 to 600 ms, 600 to 800 ms and 800 to 1000 ms).

To examine intra-individual variability of N400 effects for each EEG system, we split the data into odd and even trials. We computed the area between the related and unrelated condition curves between 300 and 800 ms, at Cz for the Neuroscan, and F3 for the EPOC+ (these electrodes were chosen as the locations where the N400 effect is likely to be prominent (Kutas & Federmeier, 2000). We used Spearman’s correlation to compare the area between the curves between odd and even trials in each individual.

#### 2.1.8 Comparison of EEG systems

We next sought to validate the EPOC+ system for recording N400 ERPs. To compare the shape of Neuroscan and EPOC+ waveforms, we ran intraclass correlations (ICC) a global index of waveforms similarities and amplitude using a previously validated Matlab script (see McArthur & Bishop, 2004),; and Spearman’s correlations, which measure the rank correlation between two waveforms, providing information about the similarity of the overall shape of the ERPs and is less sensitive to amplitude. We compared these correlations to zero by assessing whether the 95% confidence interval of the correlation values overlapped with zero, and interpreted their magnitude according to Cicchetti’s guidelines (Cicchetti, 1994). For this analysis, in order to have a fair comparison between the two systems, we re-did the pre-processing so that the Neuroscan and EPOC+ data were as comparable as possible and treated in the same way. For this, the processing proceeded as describe above except that we down-sampled Neuroscan data to match EPOC+’s sampling rate of 128 Hz and we did not remove eye-blink components from either system. We then calculated the correlations for each condition, using the entire epoch, at our two locations of interest where the electrodes from the two systems lie in close proximity: the left and right frontocentral sites (FC3 and FC4 - see locations on Figure 2). We calculated the correlation for each condition, in each individual at these two locations. We examined whether correlations were significant by computing the 95% confidence interval of the group mean and checking if they overlapped with 0 (which would correspond to no correlation). Finally, we asked whether the amplitude of the N400 effect differed between the two systems. To this end, we compared the area under the difference curve (related – unrelated ERP), using trapezoidal integration from 300 ms to 800 ms, between the two systems using a two-tailed, paired-sample t-test across individuals. These time points correspond to the expected N400 effect time course (Beukema et al., 2016; Cruse et al., 2014; Kutas & Federmeier, 2011).

**Figure 2.**
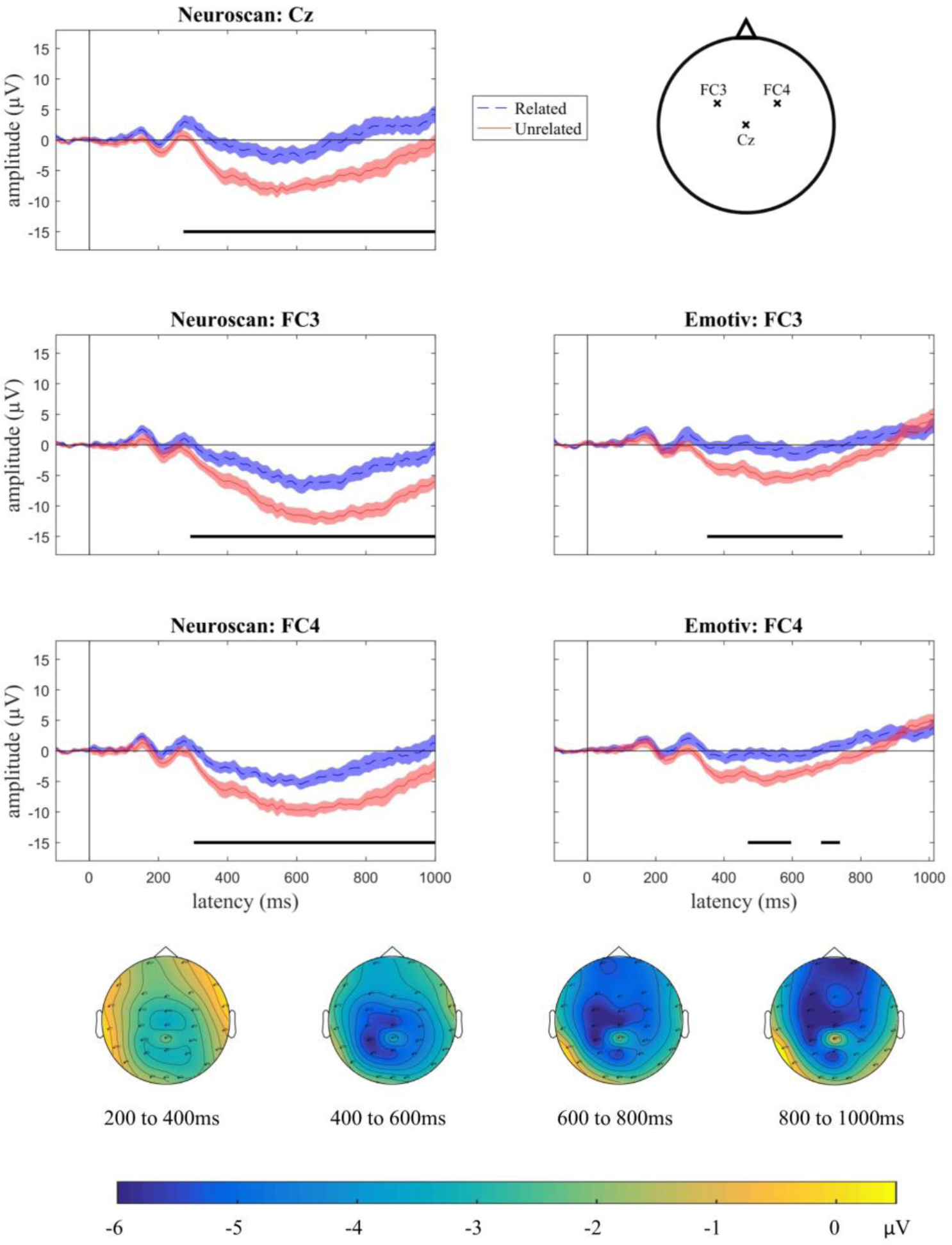
Group N400 effects for Experiment 1 (word pairs). Plots display grand average ERPs (n=16), with related (dashed blue) and unrelated (solid red) conditions for Neuroscan electrodes Cz (top left panel), FC3 (middle left) and FC4 (bottom left), and EPOC+ electrodes F3 (middle right) and F4 (bottom right). Shading indicates standard error of the mean. Time points at which there was a statistical difference between the conditions are indicated with a solid black line under the plot (p < .05, after cluster correction for multiple comparisons). Locations are shown on the top right panel. The bottom panel illustrates the topographic map of the N400 effect (unrelated minus related condition) from 200 to 1000 ms after target onset in the group for the Neuroscan system. Yellow colours indicate no difference between the two conditions and blue colours indicate a more negative-going response for the unrelated condition. The N400 effect was distributed over central and centro-frontal regions.

#### 2.1.9 Single subject temporally- and spatially-unconstrained MVPA

In order to be sensitive to individual variation in the topology and time course of N400 effects, without increasing multiple comparisons (i.e., without analysing every electrode and time point separately), we used multivariate pattern analyses (MVPA). We analysed all the data using the CosmoMVPA toolbox (Oosterhof et al., 2016) in MATLAB for Neuroscan and EPOC+ separately. First, we divided our data into a training set and a testing set, using a leave-one-target-out cross-validation approach. The training set consisted of the activation pattern across all the electrodes and time points for trials corresponding to all the targets but one, and the classifier was trained to find the decision-boundary that best distinguished between the two categories (related versus unrelated). Since each pair was repeated once, the training set consisted of 248 (62 stimuli * 2 conditions * 2 repetitions) trials. We then tested the classifier’s ability to classify the category of the remaining four trials (two related, two unrelated) corresponding to the remaining target. We repeated this procedure 63 times, each time leaving a different target out. Finally, we averaged the accuracy of the classifier for these 63 tests to yield an accuracy value for each individual.

For statistical inference, we implemented a label-permutation test in individuals (Maris & Oostenveld, 2007). For this test, we randomly permuted the condition label of the targets before classification to obtain classifier accuracies under the null-hypothesis. We performed 1000 permutations for each individual to obtain a null distribution of accuracies, to which we compared the observed (correctly labelled) accuracy. The observed accuracy was considered significantly above chance if it was larger than 95% of the accuracies in the null distribution. If the classifier performed significantly above chance, we inferred that there was information in the brain signals that differed between the two conditions (related versus unrelated words).

#### 2.1.10 Group and single-subject time-resolved MVPA

In order to examine the time course with which the brain data could be used to decode lexico-semantic condition, we ran a related, time-resolved, version of the multivariate analyses. The analyses were identical to the unconstrained MVPA, except the decoding was run for each time-point separately, using the pattern of activities from all the electrodes (rather than all time points and all electrodes in a single analysis). To test for significance we again derived an estimate of the null distribution from the data. For this, we generated 1000 random permutations of the condition labels pertaining to the different trials. For each permutation we performed the decoding analysis as before, at each time point separately. Next, we compared the observed (correctly labelled) decoding accuracy to the accuracy of these permutations. To make this comparison, we computed a threshold-free cluster enhancement (TFCE; Smith & Nichols, 2009) statistic at each time point for both the observed (correctly-labelled) data and for each of the 1000 permutations, using the cosmo_montecarlo_cluster_stat function in CoSMoMVPA.. The TFCE statistic at each time point reflects both the strength and the temporal extent of the decoding signal based on the classification accuracy at the current and neighbouring time points, allowing optimal detection of sharp accuracy peaks as well as weaker but sustained effects. We then extracted the maximum TFCE statistic across all the time points for each permutation, to create a null distribution of the maximum TFCE values over permutations (Maris & Oostenveld, 2007b). In other words, we derived a distribution of the maximum TFCE values arising across the time course by chance. Finally, we found the 95% percentile of this null distribution, and thresholded the TFCE statistic in our observed dataset with this 95% percentile value. Data at a particular time point was considered significantly above chance at the p < .05 level if its TFCE was larger than 95% of the TFCE values in the corrected null distribution. This approach corrects for multiple comparisons across time, because only those time points with TFCE values higher than the largest TFCE value across all time points in 95% of permutations are considered significant.

The group-level classifier accuracy over time was obtained by averaging each individual’s accuracy. For statistical inference at the group level, we implemented a sign-permutation test, consisting of randomly swapping the sign (positive or negative after subtracting chance, 50%) of the decoding results of each of the participants 1000 times to obtain a null-distribution of accuracies at each time point. Again, the observed accuracy and null distribution accuracies were transformed using TFCE, and accuracy was considered significantly about chance if it was larger than 95% of the TFCE values in the corrected null-distribution.

### 2.2 Results

#### 2.2.1 Behavioural results

All children had a standard score within or above the normal range (90-110) for non-verbal reasoning (K-BIT M= 123, 95% CI [114,131]) and receptive vocabulary (PPVT M= 120, 95% CI [115,126]). We asked children for a subjective rating (1-5 stars) of the performance of the aliens in each block but, as there was no ‘correct’ answer, we do not report accuracy. Children seemed to understand the instructions well and informally reported the task to be engaging.

#### 2.2.2 Group ERP analyses

At the group level, we replicated the typical N400 effect using the Neuroscan system. We found significant N400 effects at all three of our regions of interest: Cz, FC3, and FC4 (Figure 2, left panels). For the central location (Cz), the N400 effect was significant for a cluster of time points from 272 – 1000 ms, post-stimulus onset (Figure 2, top panel). For FC3, the N400 effect was significant for a cluster from 292 – 1000 ms, and for FC4 the N400 effect was significant in a cluster from 302 – 1000 ms. The group-level topographic distribution of the effect (Figure 2, bottom panel) was initially centro-parietal, spreading frontally at later time points. We were also able to record N400 effects for the group using the EPOC+ system in FC3 (from 350 to 747 ms) and FC4 (from 469 to 596, and from 684 to 739 ms), our two locations of interest (Figure 2, right panels).

#### 2.2.3 Single subject ERP analyses

Our next goal was to assess the detection rate of N400 effects in individual subjects. We defined a significant N400 effect as the presence of a statistically larger N400 in the unrelated compared to related condition (corrected for multiple comparisons across time points) in one or both of the two locations of interest that were present in both systems (FC3 or FC4). Table 1 shows the percentage of participants with significant N400 effects (“detection rate”) at each electrode. A significant N400 effect was found in 7 of the 15 (47%) participants’ Neuroscan data, and in the same number of participants’ EPOC+ data. Two participants showed an effect in the Neuroscan data but not the EPOC+ data, and vice versa. Additionally, we assessed whether inter-individual differences in the N400 effect could be explained by individual factors, such as age, vocabulary or non-verbal reasoning. We did not find any significant correlation between the amplitude of the N400, as measured by the area between the curves between 300 and 800ms recorded from Neuroscan at Cz, and age (r = -.12, p = .68), PPVT score (r = -.19, p = .52), or K-BIT score (r = -.07, p = .52).

**Table 1.** Experiment 1 (word pairs) detection rate (% of individuals) of statistically significant N400 effects in each of the three electrodes of interest, and the detection rate in a more lenient assessment where the effect was considered present if it occurred in either one or both of the two frontal electrodes.

| Electrode | FC3 | FC4 | Total (FC3 and/or FC4) | Cz |
| --- | --- | --- | --- | --- |
| Neuroscan | 47% | 33% | 47% | 47% |
| EPOC+ | 40% | 40% | 47% | N/A |

We show the individual waveforms recorded by the Neuroscan in Cz, which is where N400 effects are typically recorded (Figure 3, first and fourth columns) and in FC3, where we have both Neuroscan and EPOC+ data (Figure 3, second, third, fifth and sixth columns); FC4 results were similar. In addition, we illustrate the topography of the effect over time (Figure 4). To summarise, the detection rate of individual N400 effects was less than 50% with either EEG system, and large variations in the topography of the effect were found, suggesting inter-individual variabilities in the recorded neural signals.

**Figure 3.**
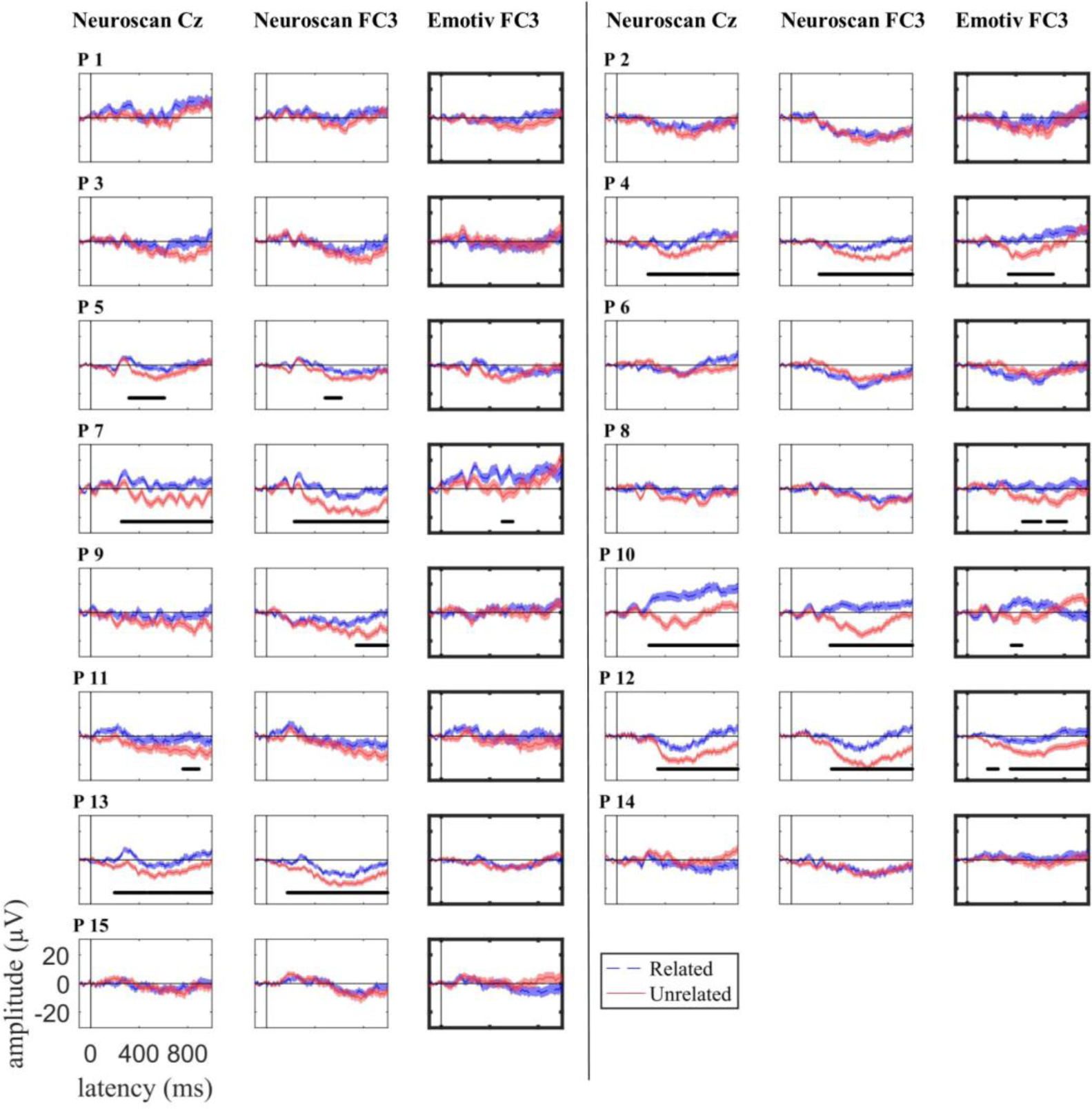
Experiment 1 (word pairs paradigm) individual participant event-related potentials to target words following a related (dashed blue) and unrelated (solid red) word for Neuroscan electrode Cz (first and fourth column), and adjacent Neuroscan and EPOC+ electrode FC3 (second, third, fifth, and sixth column), plotted ± standard error (shaded area). Time points where there was a statistically significant N400 effect in each participant and sensor are indicated with a solid, horizontal, black line. EPOC+ results are outlined in bold. P indicates participant.

**Figure 4.**
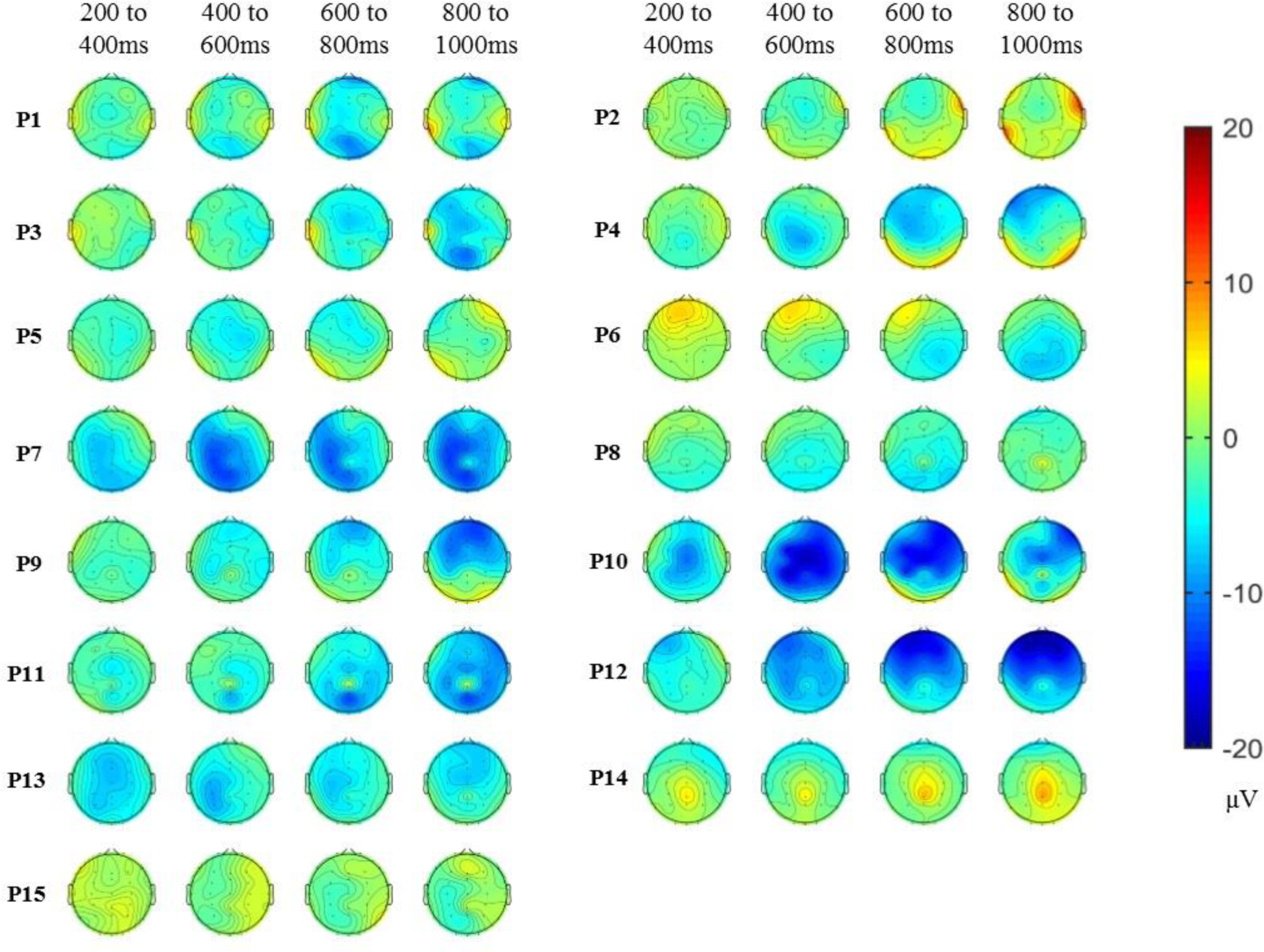
Experiment 1 (word pairs paradigm) individual topographic maps of the N400 effect (unrelated minus related condition) for 200 ms time windows from 200 to 1000 ms after target onset for Neuroscan. Red areas indicate more negative-going response for the related condition and blue areas indicate more negative-going response for the unrelated condition. The topography of the N400 effect varied across individuals. P indicates participant.

In addition, we calculated the split-half reliability of the N400 effect, by computing the Spearman’s correlation coefficient for the area under the difference curve (related – unrelated) for odds and even trials. The split-half reliability was moderate (according to the recent nomenclature by Akoglu, 2018) for Neuroscan (r = .55, p = 0.03) and weak for EPOC+ (r = .33, p = .23).

#### 2.2.4 Comparison of EEG systems

We next compared the responses of the two EEG systems using ICC and Spearman’s rank correlation (see Table 2) and by comparing the areas under the difference curves. Waveforms across the two systems were qualitatively similar in shape, and positively correlated (all Spearman’s rho ≥ 0.49, 95% CI not including zero for any of the comparisons). Mean ICC values ranged from 0.19 to 0.63 across the different conditions and sites, corresponding to fair to good correlations (Cicchetti, 1994). The ICC was significantly greater than 0 (CIs did not include 0) for the related condition on both sides and for the unrelated condition on the left side, but not for the unrelated condition on the right side. We also tested whether the amplitude of the effect was larger for Neuroscan compared to EPOC+ (as suggested by Figure 2). The area between the related and unrelated curves was numerically larger for Neuroscan than for EPOC+ both in FC3 (294 µV for Neuroscan versus 140 µV for EPOC+), and FC4 (251 µV for Neuroscan versus 82 µV for EPOC+), but the difference was not significant (FC3: t_(14)_ = 1.90, p = .0779, Cohens’ d = .46; FC4: t_(14)_ = 1.74, p = .103, Cohen’s d = .12). Therefore, the ERPs recorded by the two systems were fairly comparable in shape and not significantly different in amplitude.

**Table 2:** Experiment 1 (word pairs paradigm, bottom) mean ICC (r) and Spearman’s coeeficient (ρ) and 95% confidence intervals between waveforms simultaneously recorded with the research (Neuroscan) and gaming (EPOC+) EEG systems for the left (FC3) and right (FC4) frontocentral locations, in the semantically related and unrelated conditions. We also present the difference in area between the two condition curves between Neuroscan and EPOC+ averaged across subjects, with 95% confidence intervals.

| Condition | Coeff. | Location |  |
| --- | --- | --- | --- |
|  |  | Left frontocentral (FC3) | Right frontocentral (FC4) |
| Related | $\rho$ | 0.63 [0.51, 0.75] | 0.53 [0.37, 0.70] |
| | $r$ | 0.46 [0.31, 0.60] | 0.25 [0.03, 0.48] |
| Unrelated | $\rho$ | 0.52 [0.33, 0.72] | 0.49 [0.27, 0.71] |
| | $r$ | 0.27 [0.02, 0.52] | 0.19 [-0.1, 0.47] |
| Area between curves |  | EPOC+ : 140 [-45, 324] | EPOC+ : 82 [-113, 276] |
| (μV) |  | Neuroscan : 294 [139, 448] | Neuroscan : 251 [118, 384] |

#### 2.2.5 Decoding analyses

Our univariate N400 analyses were restricted to three *a priori* sites of interest. However, individual topography plots (Figure 4) suggested that the topography of the effect was highly variable between individuals, with effect location ranging from centroparietal to frontal sites. Therefore, we used MVPA to integrate information from across all sensors to detect differences in the neural patterns of activity to related and unrelated targets. Group level decoding performance (average over subjects) for the Neuroscan and EPOC+ data is shown in Figure 5. For Neuroscan (Figure 5, purple), classifier accuracy was statistically above chance in a cluster from 402 ms after target onset until the end of the epoch, indicating a reliable difference between the two conditions. There was no significant coding of associative context in the EPOC+ EEG data (Figure 5, yellow).

**Figure 5.**
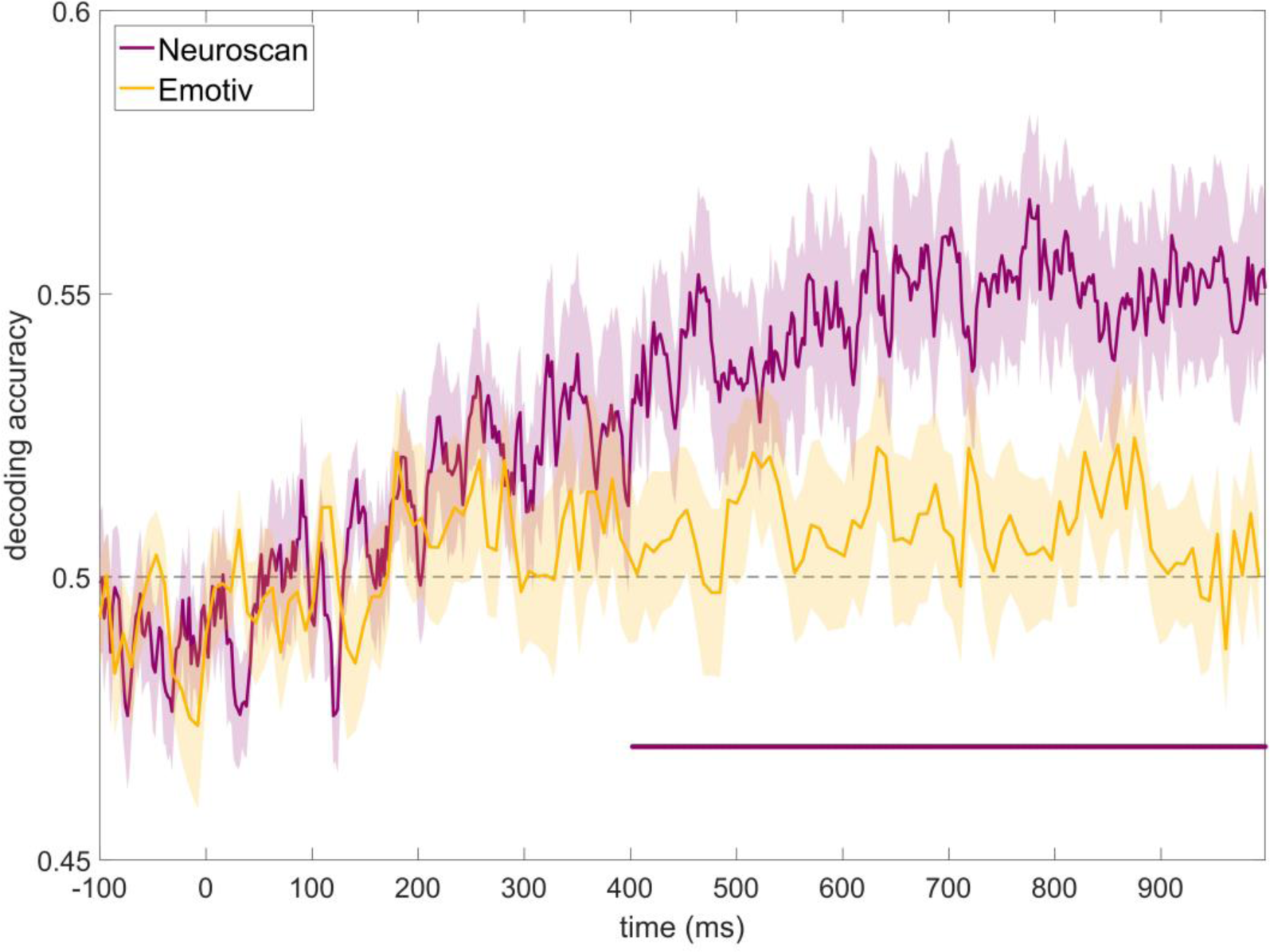
Experiment 1 (word pairs) grand average decoding accuracy for discriminating between congruent and incongruent conditions over time for Neuroscan (purple) and EPOC+ (yellow) data, shown with standard error of the mean. Time points with significant decoding for Neuroscan (p < .05, assessed with TFCE permutation tests corrected for multiple comparisons, see Methods) are shown by a purple horizontal line. Decoding accuracy was significantly above chance for Neuroscan from 402 ms but was not significant at any time point for EPOC+.

Individual decoding results for temporally- and spatially-unconstrained MVPA are shown in Figure 6. In the Neuroscan data, decoding was significant in 4/15 participants (27% detection rate). For EPOC+, the classifier only detected a significant effect in 3/15 participants (20% detection rate).

**Figure 6:**
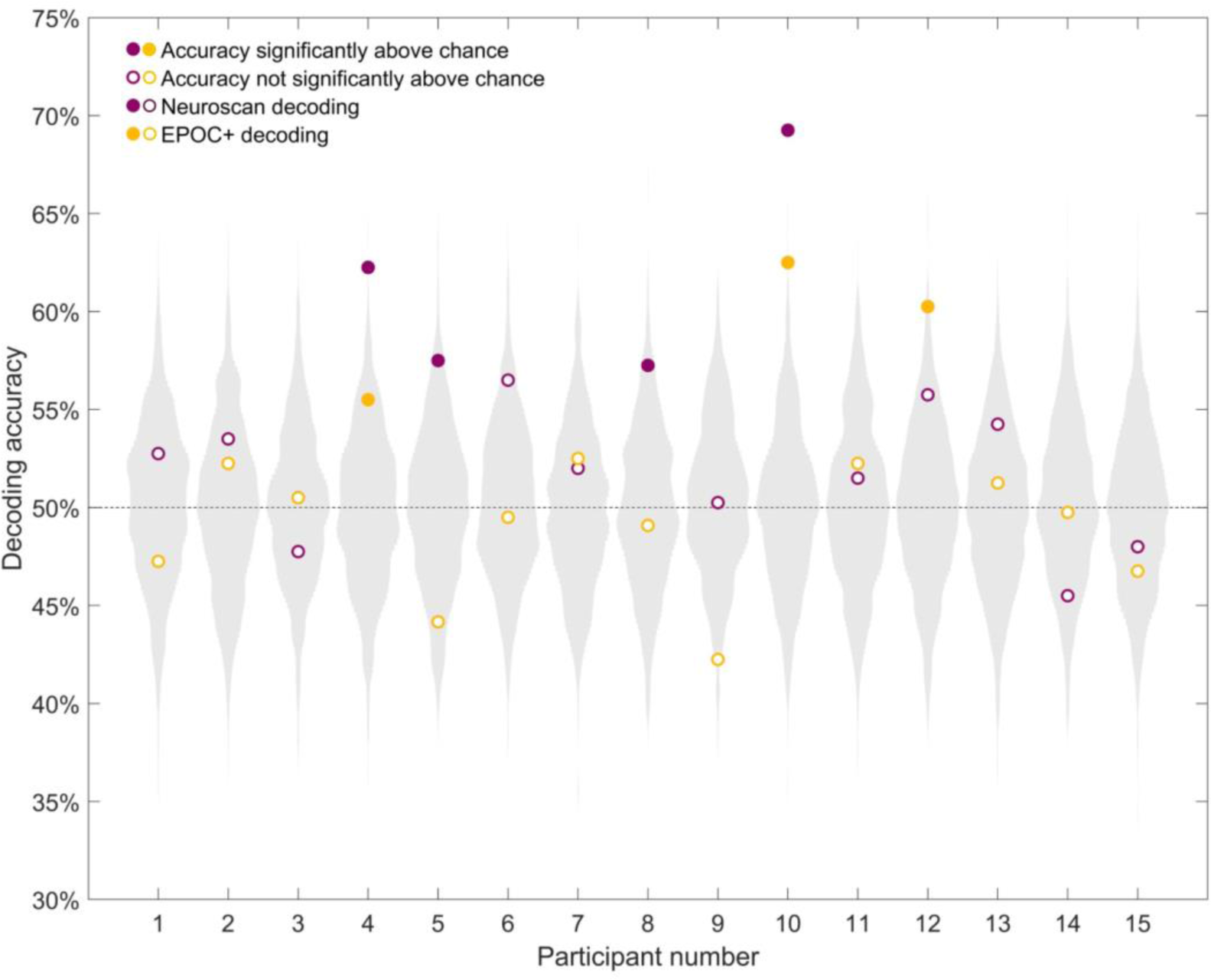
Individual decoding accuracy for classification of identical target words following related and unrelated word cues. Purple circles indicate Neuroscan decoding accuracy for each participant, while yellow circles indicate EPOC+ decoding accuracy for each participant. The grey distribution shows the null distribution obtained by the permutation test for that participant with Neuroscan (for visualisation, the null distribution for EPOC+ is not shown, as it looks similar to Neuroscan’s). Theoretical chance (50%) is indicated by the horizontal dashed line. Semantic condition could be decoded in 27% (4/15) of the participants’ Neuroscan data and in 20% (3/15) of the participants’ EPOC+ data.

In addition, we illustrated the time-course of decoding in individuals on Figure 7. For inference, we compared each individual’s decoding data to that individual’s label-permuted null distribution using a threshold-free cluster enhancement statistic (see Methods). This statistic captures both the strength of decoding and its extent in time, allowing us to conclude that even short temporal clusters reflect statistically robust differences between the conditions. When analysing decoding over time, we observe large inter-individual differences in the time course of decoding (e.g., P 14 shows significant decoding in a cluster starting at 218 ms, while P 4 shows significant decoding in a cluster starting at 608 ms). After correcting for multiple comparisons, classifier accuracy was significantly above chance in at least one temporal cluster for 53% of participants using Neuroscan data, and only 13% using EPOC+ data.

**Figure 7.**
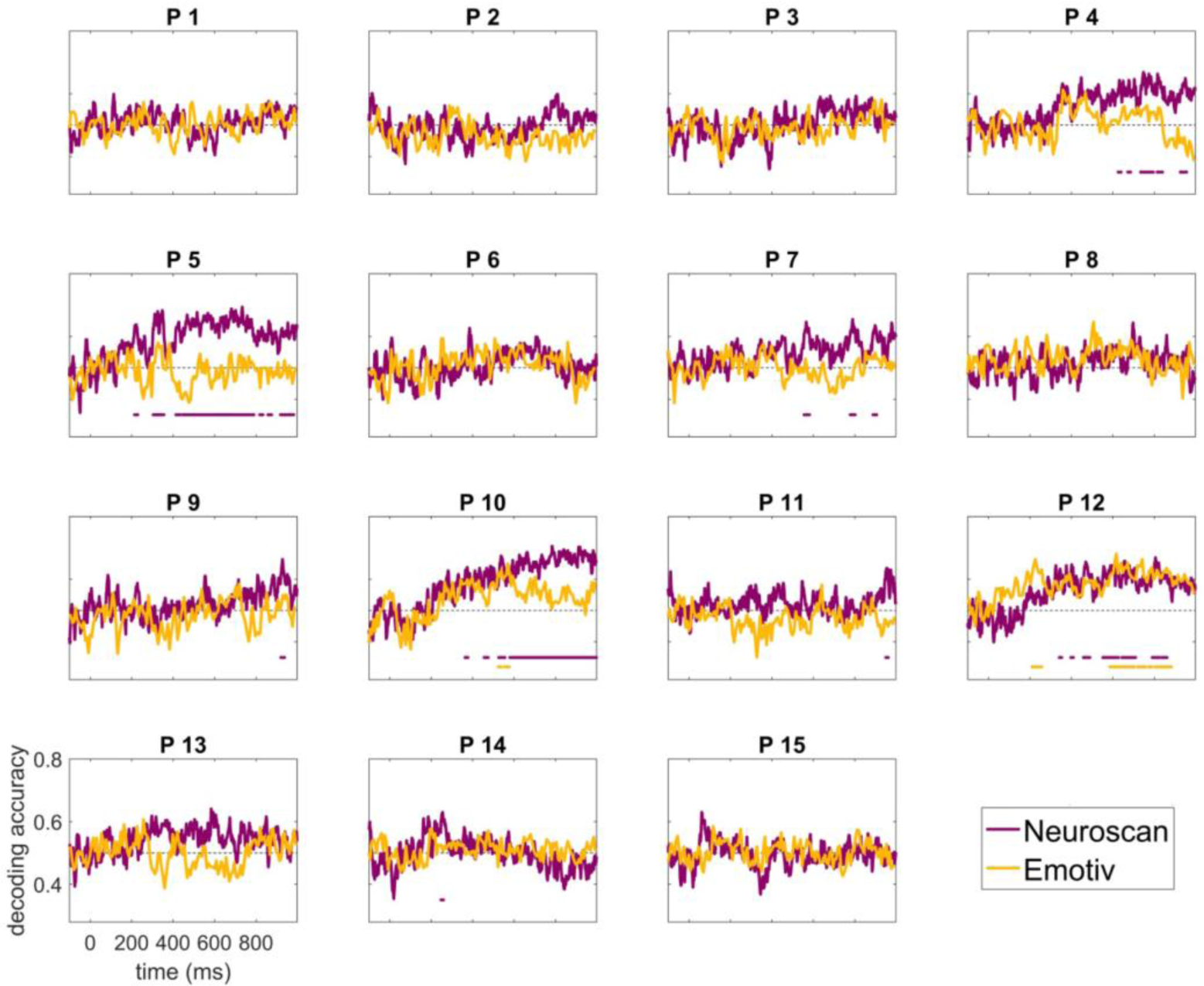
Experiment 1 (word pairs) individual participant decoding accuracy for classification of congruent *versus* incongruent conditions over time for Neuroscan (purple) and EPOC+ (yellow) data. Time points with accuracy significantly above chance (p < .05 assessed with TFCE permutation tests corrected for multiple comparisons, see Methods) are shown as solid horizontal lines (in purple for Neuroscan, and yellow for EPOC+). Theoretical chance (50%) is indicated by the horizontal dashed line. Semantic condition could be decoded in 8/15 (53%) of individuals’ Neuroscan data and in 2/15 (13%) of individual’s EPOC+ data. P indicates participant.

### 2.3 Experiment 1 summary

We examined whether differential neural responses were elicited to identical spoken words presented in different normatively-associative contexts using two different EEG systems, Neuroscan and EPOC+. We were able to elicit N400 effects in a group of children, as well as in some individual children. However, individual-subject detection rates were moderate (47% of individuals using either the Neuroscan or EPOC+ system), and the topography and timing of the effect were variable across individuals. Multivariate pattern analyses over time returned similar (53%) or weaker (13%) detection rates than traditional univariate analyses for Neuroscan and EPOC+, respectively. Waveforms for the two systems were not significantly distinguishable.

## 3 Experiment 2: congruent and incongruent sentences

As our overarching aim was to derive a sensitive measure of semantic language processing for use in individual children, we next considered another avenue to increase detection of N400 effects. A common way to elicit N400 effects is to present words in the context of semantically congruent or incongruent sentences. This may yield larger N400 effects than the word pairs task, as sentences provide a stronger semantic context compared to a single probe word (Kutas, 1993). In addition, any deleterious effect of repeating stimuli (which is necessary to perfectly match the stimuli across conditions) may be attenuated in sentences since so many words are presented on each trial (e.g., Cruse et al. 2014). For this reason, our second experiment used words presented in sentences. We also modified the task that participants performed to make it more demanding and encourage greater attention to the stimuli than for the word pairs task.

### 3.1 Methods

#### 3.1.1 Participants

Eighteen participants, aged 6 to 12 years, were recruited as described for Experiment 1. The data from two participants were excluded due to excessive artefacts in the EEG data. The final set of data thus came from 16 participants (mean age =10.3, SD=2.4, 8 male and 8 female), five of whom had also participated in Experiment 1.

#### 3.1.2 Stimuli

We created two conditions: (1) *congruent sentences*, which were semantically correct (e.g., “she wore a necklace around her *neck*”); and (2) *incongruent sentences*, which ended with an anomalous word (e.g., “There were candles on the birthday *neck*”). The set of congruent sentences were based on 56 high-close probability sentences from the norms of Block and Baldwin (2010) and were chosen according to suitability for children and such that target words were high-frequency words acquired by age five (Kuperman et al., 2012). We recombined sentence stems and target words to form the set of incongruent sentences. We ensured that the incongruent target word was unexpected but grammatically correct. It also did not begin with the same phoneme or rhyme with the corresponding congruent target word. Within one session, each sentence stem and target was used twice, once in the congruent condition and once in the incongruent condition. The final set of stimuli consisted of 56 sentences in each condition, 112 in total (see supplementary table S2 for the complete list).

The recorded stimuli were provided by a collaborator in England (S.Y.). The sentences were digitally recorded by a female native British English speaker in a soundproof room and edited in Audacity®. To avoid co-articulation, the speaker recorded the sentence stems separately from the target words. This also introduced a lengthening in the final word of the sentence stem. Sentence stems and targets were combined online during stimulus presentation with a 100 ms silence between the sentence frame and the target word. The target words had a mean length of 464 ms (SD: 89 ms, range: 309-700 ms).

#### 3.1.3 EEG Equipment

The equipment and experimental setup were the same as in Experiment 1, including the completion of the matrices section of the Kaufman Brief Intelligence Test, Second Edition (K-BIT), and the Peabody Picture Vocabulary Test, Fourth Edition (PPVT). Participants who completed Experiment 1 did not complete these tests a second time.

#### 3.1.4 Experimental procedure

Participants completed two EEG recording sessions of 25 minutes, separated by a 5-minute break. Each session included all 112 sentences (56 congruent, 56 incongruent). We presented the sentences in a pseudo-random order that was reversed in the second session. We optimised the order to avoid bias in the sequence of related and unrelated trials as described above, with all sentences presented once before being repeated in the alternate condition, and to maximise the distance between repeated presentations of the same target word. We allocated the sentences to this trial order pseudo-randomly with the additional constraint that there were at least two sentences between any repetitions of semantic content in the sentence frame or target word. We presented an image of a satellite centrally on the screen to signal the onset of each trial, and kept this display on for the whole trial. This served as an alerting cue and encouraged children to fixate, reducing eye-movements. After 2 s, we presented the sentence through the speakers and the satellite remained onscreen for a further 1.5 s after the presentation of the target word. There was then a 2 s inter-trial-interval before the next trial. Each 20-minute session consisted of 16 blocks of 4 to 10 trials, after which children gave an answer to the experimenter (see below).

We designed a task that was strongly engaging for children while requiring minimal overt responses. It was embedded in the context of a story: an evil alien Lord had messed up some of the “messages” that we were trying to send to our extra-terrestrial friends. Participants were asked to pay attention to each sentence and to count how many did not make sense. Accurate responses, given at the end of each block, would help “catch” an evil alien’s henchman who appeared on the screen. This encouraged participants to pay attention and to make covert semantic judgments of sentences. Most of the children appeared to be highly engaged and motivated by the task, and reported that they found it entertaining. The whole experiment, including setup, took approximately 2 hours.

#### 3.1.5 Offline EEG processing, ERP, and MVPA

The correlation scores, ERP analyses, and decoding analyses were performed as for Experiment 1.

For the Neuroscan data, an average of 12 epochs (11%) for the related condition (SD = 8.38) and 13 epochs (12%) for the unrelated condition (SD = 8.27) were rejected. For the EPOC+ data, an average of 8 epochs (7%) for the related condition (SD = 6.88) and 6 epochs (5%) for the unrelated condition (SD = 5.14) were rejected.

### 3.2 Results

#### 3.2.1 Behavioural results

All participants scored within or above the normal range for non-verbal reasoning (K-BIT score M= 111, 95% CI [101,120]) and receptive vocabulary (PPVT score M=117, 95% CI [111,122]).

Participants performed the behavioural task with a high degree of accuracy (mean percent correct: M=96.29%, SD=3.36%, range = [88.4%, 100%]), indicating that they understood the sentences, and were able to notice semantic anomalies.

#### 3.2.2 Group ERP analyses

We recorded large N400 effects in the group using Neuroscan in all of the electrodes of interest (Figure 8, left panels). We also recorded N400-like effects using the EPOC+ system at our two locations of interest, FC3 and FC4. However, these effects only reached significance at FC4, possibly indicating a lesser sensitivity of the EPOC+ system (Figure 8, right panels).

**Figure 8.**
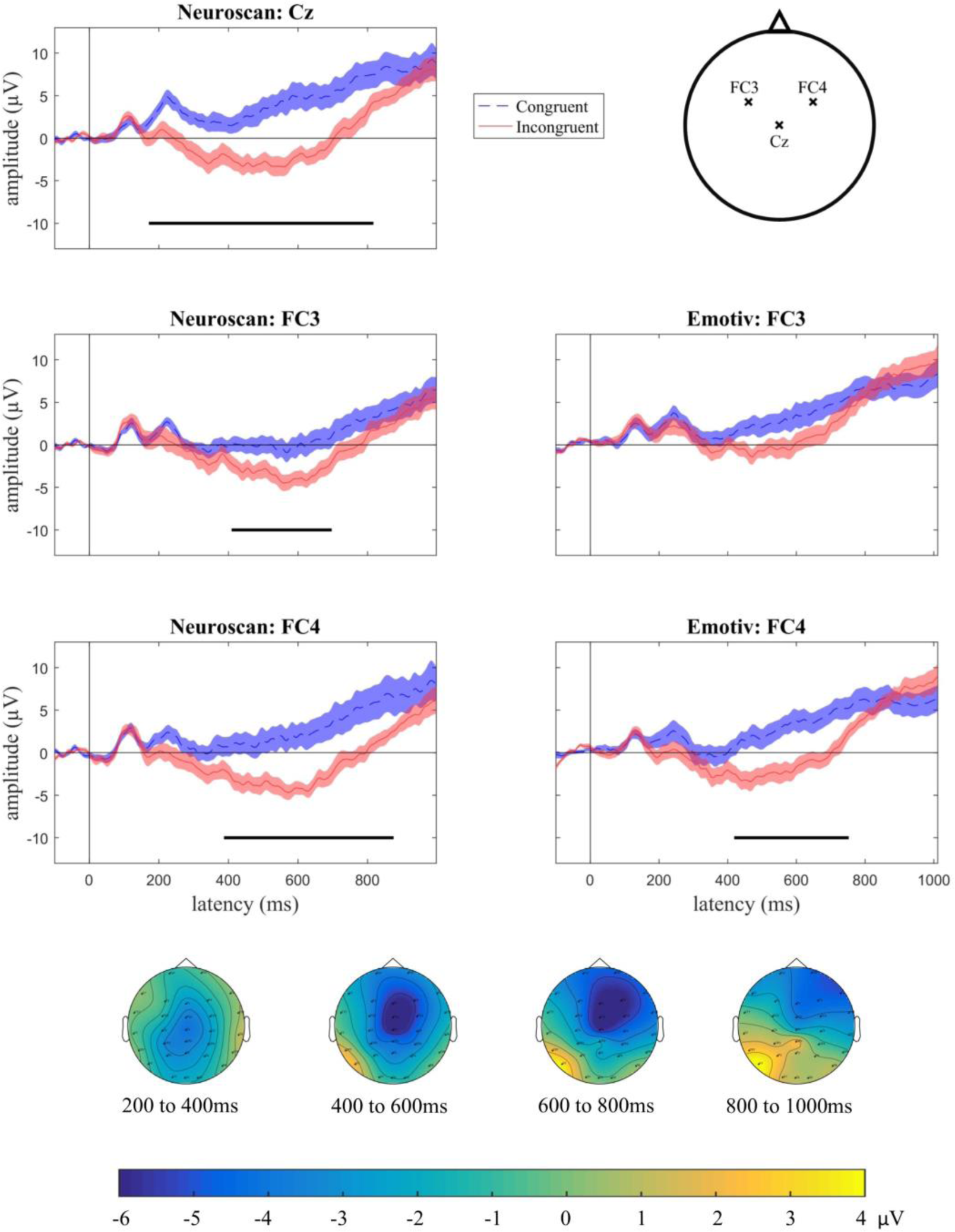
Group N400 effects for Experiment 2 (sentence paradigm). Plots show grand average ERPs (n=16), with congruent (dashed blue) and incongruent (solid red) conditions for Neuroscan electrodes Cz (top left panel), FC3 (middle left) and FC4 (bottom left), and EPOC+ electrodes F3 (middle right) and F4 (bottom right). Shading indicates standard error of the mean. Temporal clusters at which there was a statistical difference between the conditions are indicated with a solid black line (p < .05, after cluster correction for multiple comparisons). The bottom panel illustrates the topographic map of the N400 effect (incongruent minus congruent condition) from 200 to 1000 ms after target onset in the group (n=16). Yellow areas indicate a more negative-going response for the congruent condition and blue areas indicate a more negative-going response for the incongruent condition. The N400 effect was mainly distributed over the centro-frontal region.

For the central location (Cz), the N400 effect started at 171 ms and continued until 817 ms post-stimulus onset (Figure 8, top panel). For the frontal sites, the effect started later with a significant cluster from 409 – 697 ms for FC3 (Figure 8, middle left panel) and a significant cluster from 387 – 875 ms for FC4 (Figure 8, bottom left panel). For EPOC+, the N400 effect was significant in FC4 for a cluster from 418 – 752 ms. Potentials in both conditions and all three sensors also shifted in the positive direction over time, possibly corresponding to the closure positive shift, an ERP component reflecting the processing that occurs at a prosodic boundary (Steinhauer et al., 1999; Steinhauer & Friederici, 2001).

The topographic distribution of the N400 effect is shown in Figure 8, bottom panel. The distribution was centro-frontal with a slight right bias. This is perhaps in line with previous reports that found the N400 effect to be more frontal in children than in adults (e.g., Friedrich & Friederici, 2004; Henderson et al., 2011a found a centro-frontal distribution of N400 effects for infants and children, albeit using a picture-word paradigm), although we did not observe this in Experiment 1.

#### 3.2.3 Single subject ERP analyses

We then assessed the detection rate of neural responses in individual subjects. We observed reliable N400 effects in one or both of FC3 and FC4 in 56% of the participants with the Neuroscan data and 50% in the EPOC+ data (detection rates are shown in Table 3). We again did not find any significant correlation between the amplitude of the N400 effect and age (r = -.13, p = .63), PPVT score (r = .25, p = .37), or K-BIT score (r = .41, p = .37). The topography of the effect was again variable between participants, ranging from frontal (P1, P11, P14) to parietal locations (P13).

**Table 3:** Experiment 2 (sentence paradigm) detection rate (% of individuals) of statistically significant N400 effects in each of the three electrodes of interest, and the detection rate in a more lenient assessment where the effect was considered present if it occurred in either one or both of the two frontal electrodes.

| Electrode | FC3 | FC4 | Total (FC3 and/or FC4) | Cz |
| --- | --- | --- | --- | --- |
| Neuroscan | 38% | 50% | 56% | 50% |
| EPOC+ | 38% | 44% | 50% | N/A |

Figure 9 shows individual participant waveforms for Neuroscan electrode Cz (Figure 9, first and fourth column), and for Neuroscan (Figure 9, second and fifth column) and EPOC+ electrode FC3 (Figure 9, third and sixth column), which was the site with the highest detection rate. We again found that the topographical distribution of the effect was variable across individuals (Figure 10).

**Figure 9.**
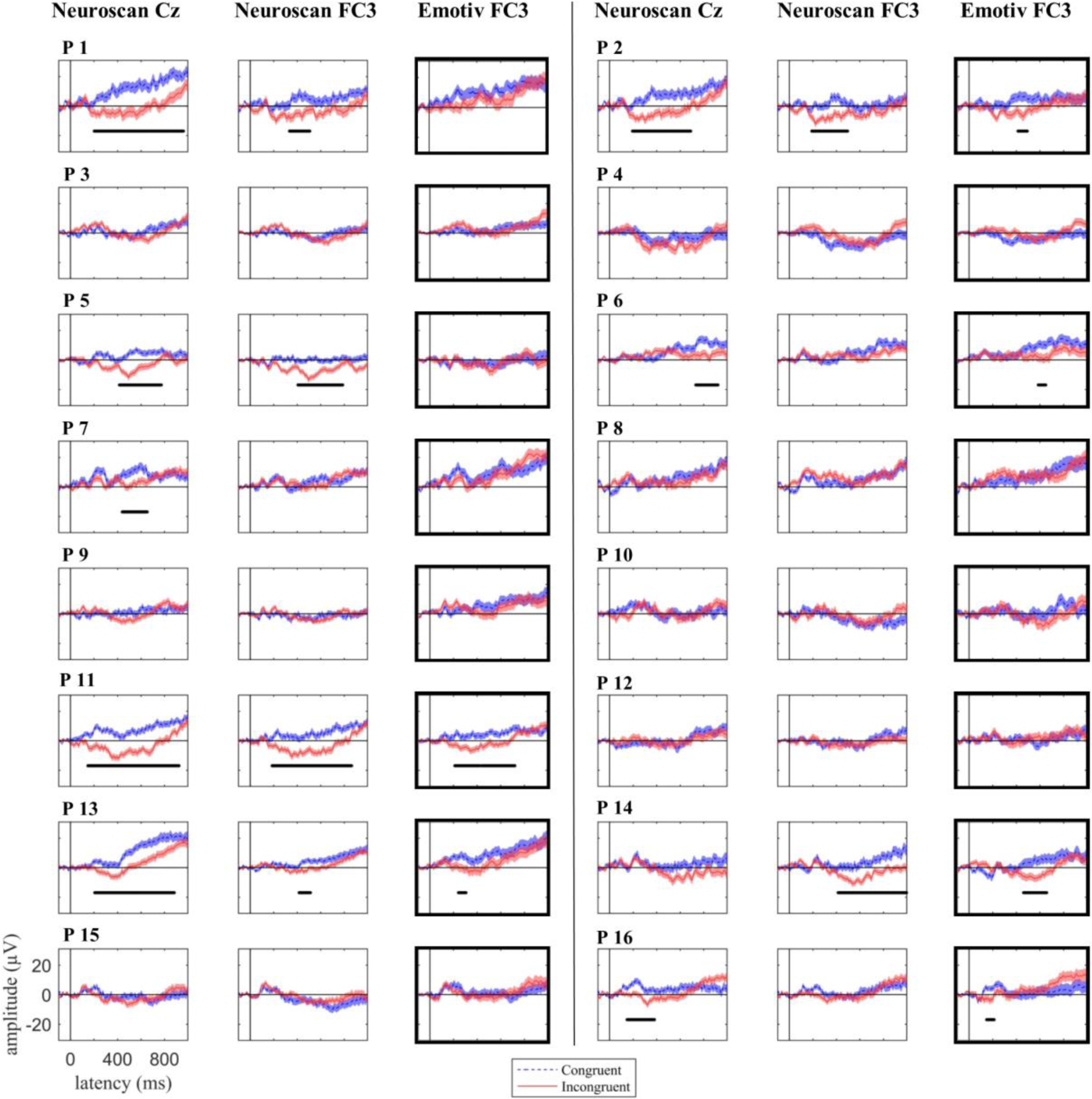
Experiment 2 (sentence paradigm) individual responses to target words in congruent (dashed blue) and incongruent (solid red) sentences for Neuroscan electrode Cz (first and fourth column), and adjacent Neuroscan and EPOC+ electrode FC4, plotted ± standard error (second, third, fifth, and sixth column). Statistical N400 effect is shown as a solid black line. EPOC+ results are outlined in bold. P indicates participant.

**Figure 10.**
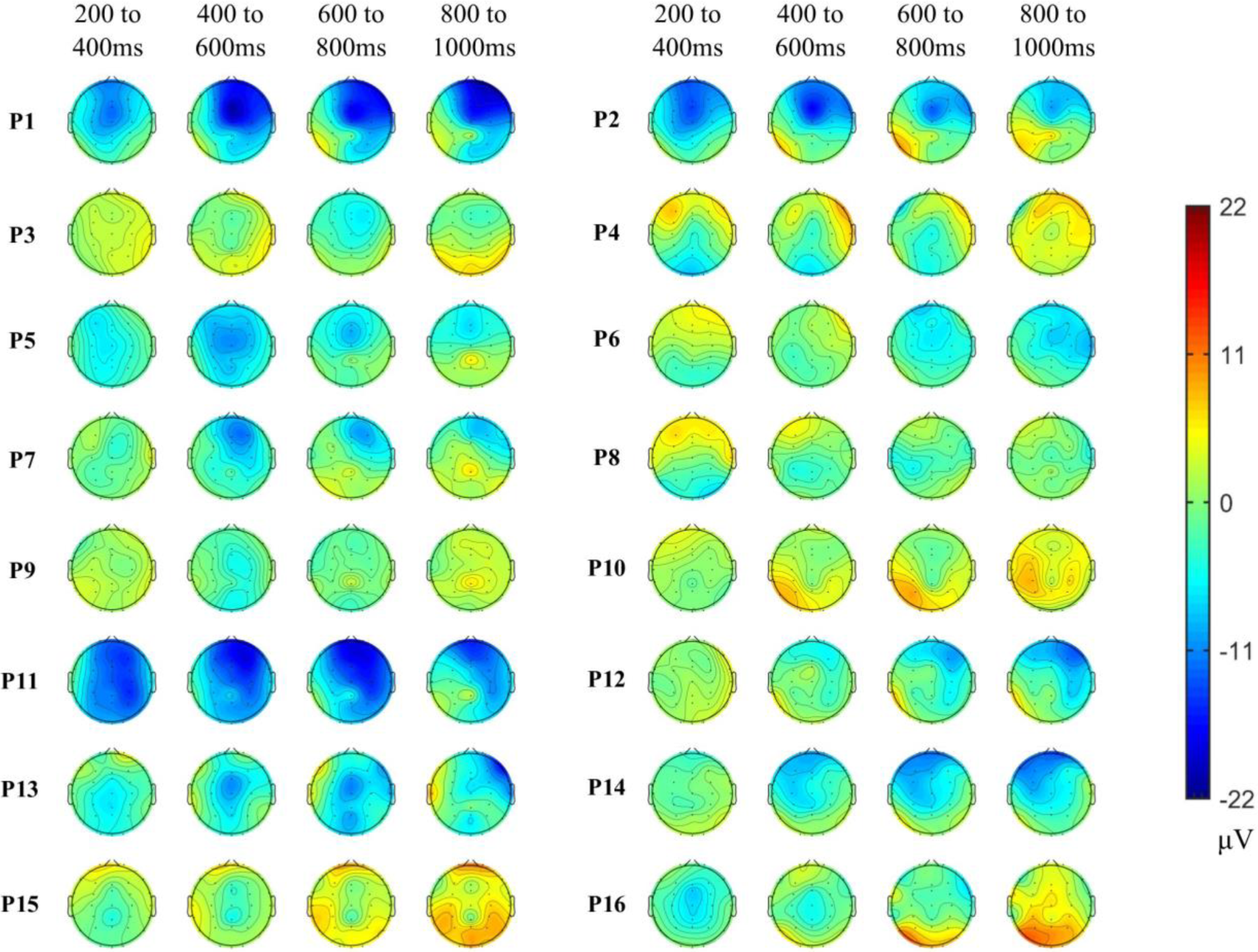
Experiment 2 (sentence paradigm) individual participant topographic maps of the N400 effect (incongruent minus congruent condition) for 200 ms time windows from 200 to 1000 ms after target onset for Neuroscan. Red areas indicate more negative-going response for the congruent condition and blue areas indicate more negative-going response for the incongruent condition. The N400 effect location was variable across individuals. P indicates participant.

Again, we calculated the split-half reliability of the N400 effect across individuals by computing the Spearman’s correlation coefficient for the area under the difference curve (related – unrelated) for odds and even trials. The split-half reliability was again moderate for Neuroscan (r = .56, p = 0.02). For EPOC+, the reliability was null (r = -.04, p = .87), potentially reflecting poor data quality with this system when analysing only half of the trials.

#### 3.2.4 Comparison of EEG systems

We again compared the responses of the two systems directly using ICC and Spearman’s rank correlation and by comparing the area under the difference curve at our two locations of interest (see Table 4). Waveforms for the sentences task across the two systems were qualitatively fairly similar in shape and amplitude, as indicated by significant positive correlations (ICC ≥ 0.23 CI not including zero for any of the comparisons) for both sites and conditions. The area between the related and unrelated curves was numerically larger for Neuroscan than for EPOC+ both in FC3 (208 µV for Neuroscan versus 125 µV for EPOC+), and FC4 (340 µV for Neuroscan EPOC+ versus 196 µV for EPOC+), but the difference was not significant for either site (FC3: t_(15)_ = 0.82, p = .43, Cohen’s d = .71 FC4: t_(15)_ = 1.44, p = .17, Cohen’s d = .65).

**Table 4:** Experiment 2 (sentence paradigm) mean ICC (r) and Spearman’s coeeficient (ρ) and 95% confidence intervals between waveforms simultaneously recorded with the research (Neuroscan) and gaming (EPOC+) EEG systems for the left (FC3) and right (FC4) frontocentral locations, in the semantically congruent and incongruent conditions. We also present the difference in area between the two conditions for Neuroscan and EPOC+ averaged across subjects with 95% confidence intervals.

| Condition | Coeff. | Location |  |
| --- | --- | --- | --- |
|  |  | Left frontocentral (FC3) | Right frontocentral (FC4) |
| Congruent | $\rho$ | 0.57 [0.38, 0.76] | 0.54 [0.32, 0.77] |
| | $r$ | 0.23 [0.01, 0.46] | 0.30 [0.02, 0.58] |
| Incongruent | $\rho$ | 0.71 [0.58, 0.85] | 0.72 [0.58, 0.87] |
| | $r$ | 0.53 [0.34, 0.72] | 0.62 [0.44, 0.80] |
| Area between curves |  | EPOC+ : 125 [-39, 290] | EPOC+ : 196 [52, 339] |
| (μV) |  | Neuroscan : 208 [42, 373] | Neuroscan : 340 [138, 541] |

#### 3.2.5 Decoding analyses

We again tested whether our detection rate would improve by combining data across sensors using MVPA. At the group level, we saw significant decoding of semantic context (congruent or incongruent sentence frames) in the Neuroscan data from 162 ms to 1000 ms (Figure 11, purple line). We also decoded the semantic category from the EPOC+ group level data (Figure 10, yellow line), at several clusters of time points between 414 ms and 860 ms. Therefore, MVPA decoding matched the time course of univariate decoding seen at the group level in Neuroscan and EPOC+ data (Figure 8).

**Figure 11.**
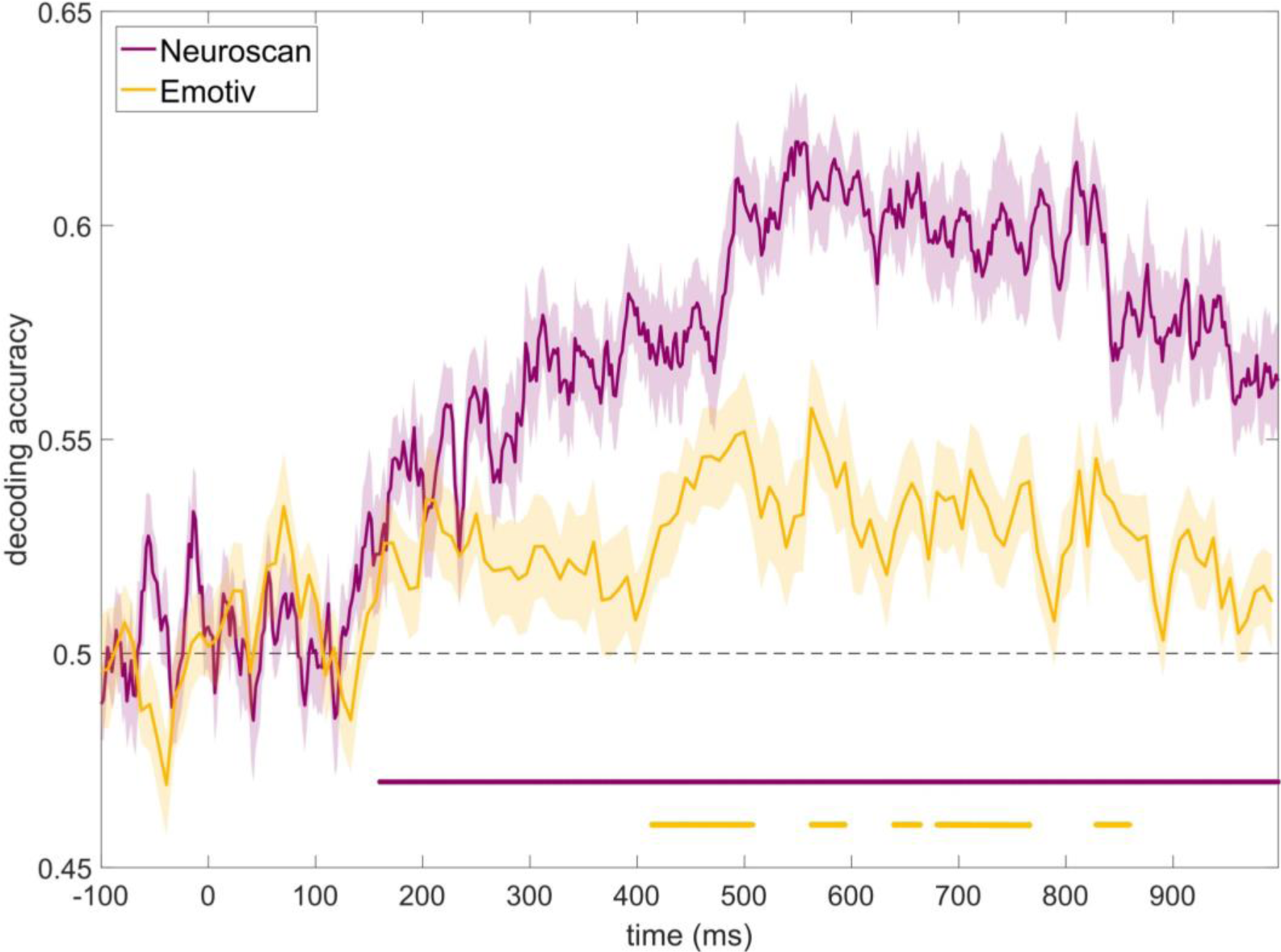
Experiment 2 (sentence paradigm) grand average decoding accuracy for discriminating between identical target words presented in congruent and incongruent conditions at each time point for Neuroscan (purple) and EPOC+ (yellow) data. Shading indicates standard error of the mean. Clusters of significant decoding are shown by a purple (Neuroscan) and yellow (EPOC+) horizontal line. Decoding accuracy was significantly above chance for Neuroscan in a cluster from 162 ms to 1000ms, and for EPOC+ at several clusters of time points between 414 ms and 860 ms.

At the individual level, using temporally- and spatially-unconstrained MVPA, we could decode the semantic condition from the Neuroscan data in all but two participants (88% detection, Figure 12, purple circles). This was a marked improvement relative to the univariate detection rate at individual channels (38-50%, above). However, for EPOC+, the classifier only detected a significant effect in 6 of the 16 participants (38% detection rate, Figure 12, yellow circles).

**Figure 12:**
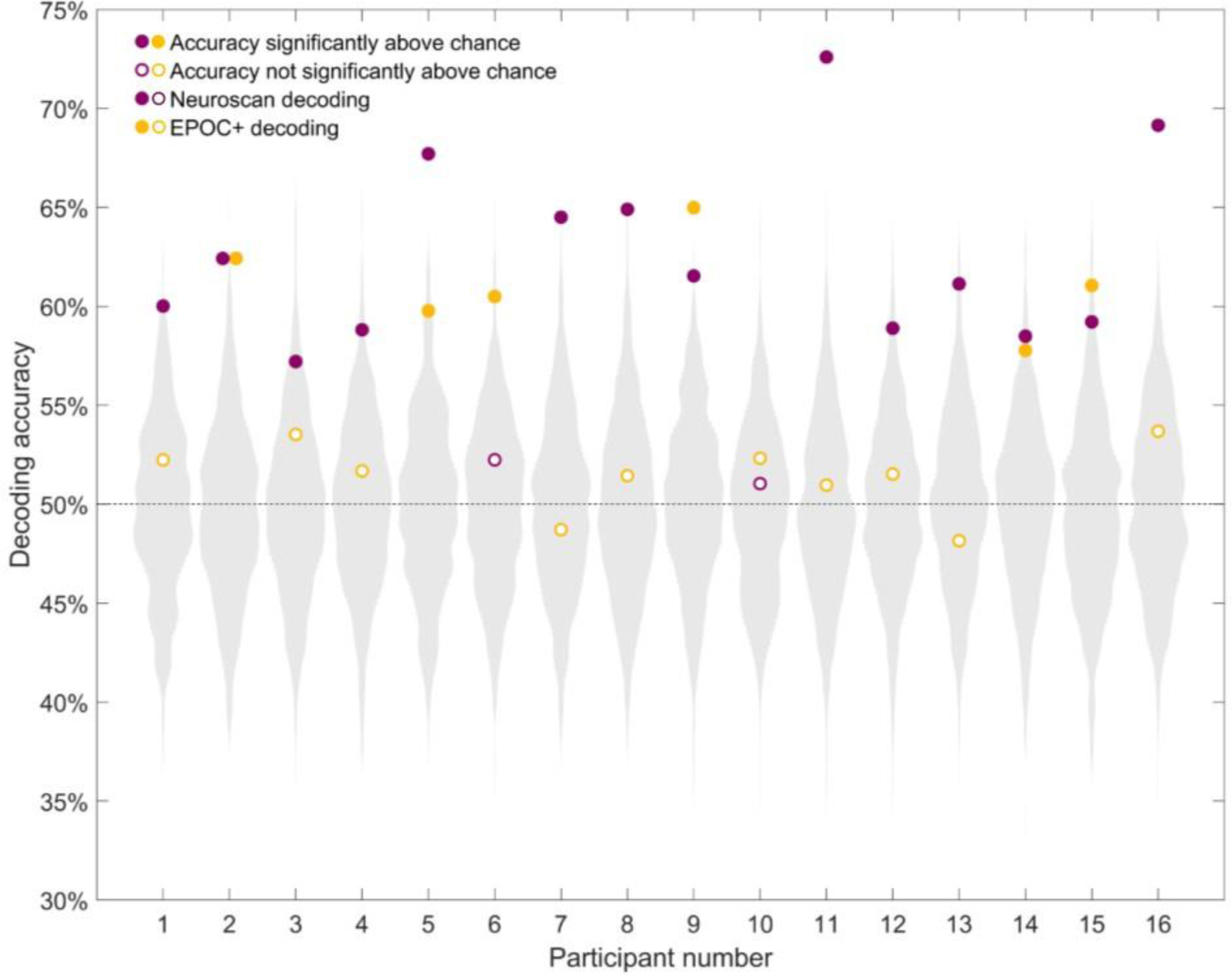
Individual decoding accuracy for classification of identical target words in congruent and incongruent contexts. Purple circles indicate Neuroscan decoding accuracy for each participant, while yellow circles indicate EPOC+ decoding accuracy for each participant. The grey distribution shows the null distribution obtained by the permutation test for that participant with Neuroscan (for visualisation, the null distribution for EPOC+ is not shown, in practice it looks similar to Neuroscan’s). Theoretical chance (50%) is indicated by the horizontal dashed line. Semantic condition could be decoded in 88% (14/16) of the participants’ Neuroscan data and in 38% (6/16) of the participants’ EPOC+ data.

In addition, we illustrate the time course of decoding, by using MVPA over time. After correction for multiple comparisons, we detected at least one temporal cluster in 88% of participants with the Neuroscan data (Figure 13, purple traces), but in only 25% of participants for EPOC+ (Figure 13, yellow traces).

**Figure 13.**
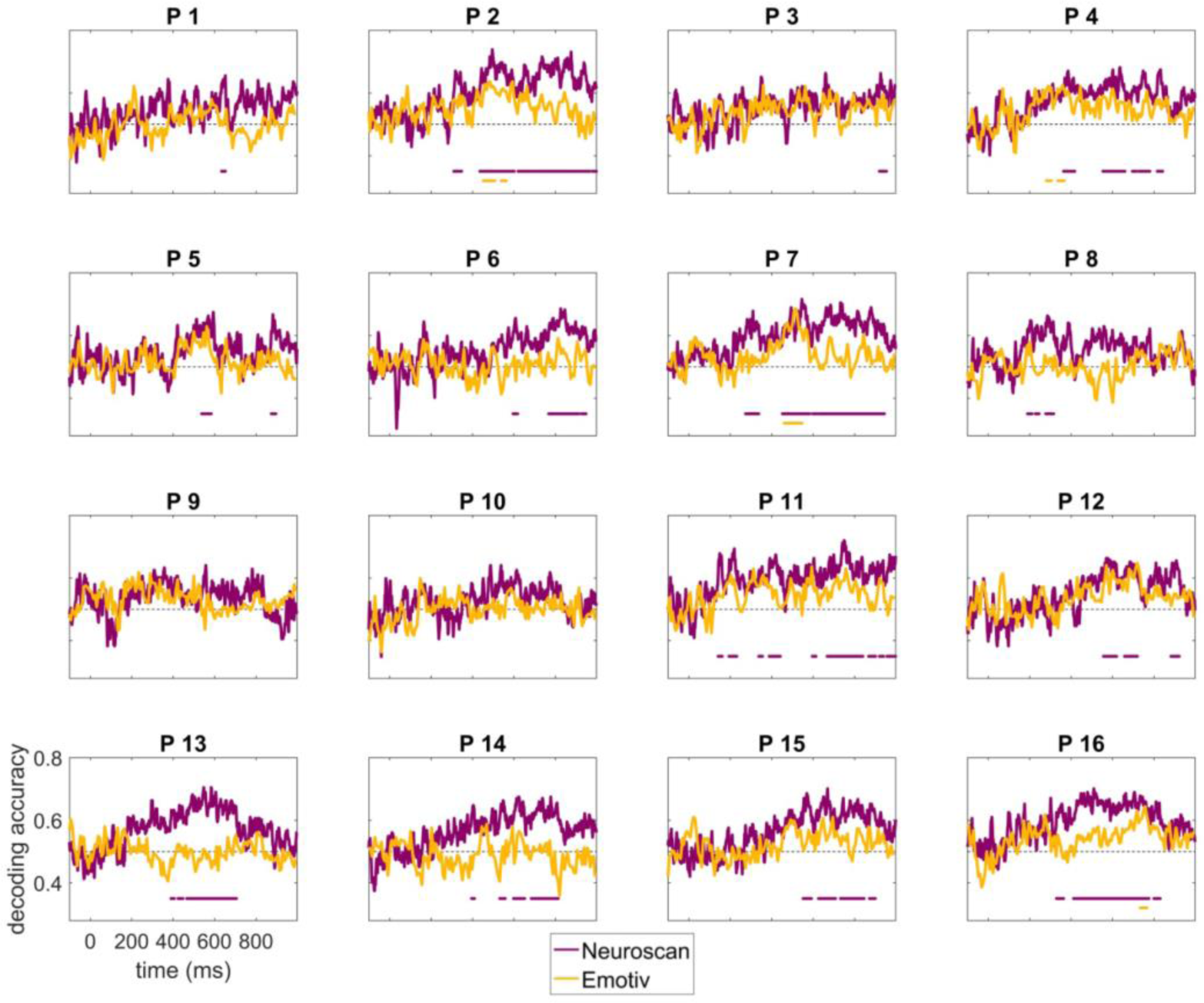
Experiment 2 (sentence paradigm) individual decoding accuracy for discriminating between congruent and incongruent conditions over time for Neuroscan (purple) and EPOC+ (yellow) data. Time points with accuracy significantly above chance (temporal cluster correction, p < .05) are shown as solid horizontal lines (in purple for Neuroscan, and yellow for EPOC+). Semantic condition could be decoded in 88% of the participants’ Neuroscan data and in 25% of the participants’ EPOC+ data. P indicates participant.

Finally, we examined whether the superior decoding of Neuroscan could be attributed to the larger number of electrodes (33 in Neuroscan versus 12 in EPOC+) by performing an additional time-resolved MVPA analysis on the Neuroscan data using only the 12 electrodes closest to the EPOC+ electrodes. In these conditions, the MVPA again performed well for the Neuroscan data, identifying statistical differences in the signal for 88% of participants. Thus, the Neuroscan data appeared to be more suitable for decoding than the EPOC+ data and the difference could not be attributed to the difference in the number or location of electrodes.

### 3.3 Experiment 2 summary

Using a paradigm that contrasted congruent and incongruent sentences, we elicited a univariate N400 effect in a group of children, and in up to 50% of the participants using Neuroscan and 44% using EPOC+. Using multivariate analyses, we decoded the semantic condition in all but two individuals (88%) using the Neuroscan data. This suggests that multivariate analyses, which take the pattern across all electrodes into account, may be a sensitive way to detect effects in individuals, accounting for the topographic variability of effects across individuals. However, decoding was, if anything, slightly worse than univariate analyses for the EPOC+ system (25% with MVPA, as opposed to 44% with univariate analyses), suggesting that MVPA may be more sensitive to the differences in data quality from research-grade and gaming EEG systems than traditional analyses.

## 4 General Discussion

For children with impaired communication, objective and reliable tests of language comprehension are urgently needed. Here, we examined the consistency of neural signals across neurotypical children using two auditory N400 paradigms. We assessed the detection of discriminative brain signals in response to identical spoken words presented in the context of a congruent and incongruent single word prime (Experiment 1) or sentence frame (Experiment 2), and compared the signal recorded by a low-cost gaming EEG system, Emotiv EPOC+, to that recorded by a traditional research EEG system, Neuroscan SynAmps2. We further used two approaches to evaluate the electrophysiological response: univariate analysis of the N400 effect, and multivariate decoding. We found that the N400 effect could be observed at the group level, and detected at an individual level in about half of the children, using either paradigm (word pairs or sentences) and using either EEG system. However, the sentences paradigm, in which children counted the incongruent sentences, was more promising, with the best individual participant detection rate, 88%, given by multivariate analyses of the Neuroscan data.

Despite an extensive body of literature on the N400 effect, only a few studies have carried out statistical testing at the individual subject level or report individual participants’ waveforms. In our data, statistically significant traditional N400 effects were present in about half of our participants using either normatively-associated word pairs or sentences to induce a violation of lexical-semantic predictions. This may seem low, given that the N400 is widely reported to be a large and robust effect (e.g., Kutas & Federmeier, 2011), but it is in fact similar to statistical detection rates in adults using similar tasks (Cruse et al., 2014). This raises an important question about how robust and prevalent individual N400 effects actually are. At a minimum, it points to large inter-individual variability in presence and strength of neural responses. There are several possible explanations for this variability.

It is possible that we simply lacked sensitivity to detect consistent neural effects within testing constraints (i.e., we needed more data to detect an effect in every individual). This is difficult to quantify because traditional power analyses fail to capture the cluster-based multiple-comparisons correction we used in our univariate analyses. We do note however that the signal was noisy at the individual-level, as illustrated by individuals’ ERP waveforms. This is due to averaging fewer trials together compared to grand average ERPs, and the possibility that trials are noisier than in adults to begin with because of children’s movements during the task. This low signal-to-noise ratio is reflected by medium to low split-half intra-individual reliability of the N400 effects (r ≈ .5 for Neuroscan in both experiments; r = .33 and r = -.04 for EPOC+ in each experiment respectively). This is lower than previously reported N400 reliability data (e.g., Kiang et al. (2013) reported high reliability (r = .85) in typical adults with an average of approximately 150 usable trials).

It is also possible that lexico-semantic differences and incongruencies were not processed similarly at a cognitive level across individuals. For example, individuals may vary in the extent to which they attempt to integrate the unpredicted target word into the semantic context or process differently the wider experiment probabilistic design, where predictions were violated 50% of the time. In addition, or instead, it is possible that individuals may have similar cognitive processing but the neural substrates supporting this processing vary. Previous studies have reported that N400 effects vary with age (e.g., Atchley et al., 2006; Holcomb et al., 1992; Juottonen et al., 1996) non-verbal intelligence (Jaušovec & Jaušovec, 2000) and vocabulary knowledge (Byrne et al 1995, although see Henderson et al 2011b for an absence of relationship with vocabulary knowledge). In our data, there was no simple explanation for inter-individual variability in N400 - we did not find any association of N400 with vocabulary or non-verbal intelligence score. However, since our sample size was very small (n = 15 and 16) and our stimuli were not designed to test these associations, we do not draw any strong conclusion from this.

The details of the stimuli and the participants’ task also seem likely to affect the size of the N400 (e.g., Bentin et al., 1993; Chwilla et al., 1995; Ortega et al., 2008; Perrin & García-Larrea, 2003) and therefore, presumably, individual-subject detection rate. For example, Cruse et al. (2014) reported superior individual-subject detection rates for active and overt tasks relative to passive tasks. Cruse et al. also reported superior rates for normatively associated word pairs (50%) than for sentences (17%) when heard passively. However, we found that the response to the two stimulus sets were comparable, and, if anything, slightly better for sentences, when we considered an effect recorded from either of our sites of interest (47% in word pairs, and 56% in sentences). The numerically lower sensitivity of our word pair paradigm compared to the sentence paradigm may be due to the repetition of the target words during the experiment (each target word was repeated between two and four times per condition). Stimuli repetition has been found to reduce the strength of the N400 effect, especially in word pair paradigms (Duncan et al., 2009, Cruse et al., 2014), but may be less of a problem when using sentence stimuli, as they contain many words which are naturally repeated (Cruse et al., 2014).

Our individual-participant data also emphasise the variability in topography and time course of individual N400 responses (previously suggested by Henderson et al., 2011)), which may be particularly important to consider when testing children and special populations, and might necessitate different analysis approaches. For example, in those participants not showing a reliable N400 effect, it is possible that univariate analyses failed to detect more subtle changes in the neural responses or that N400-like responses occurred in EEG channels that were not analysed. To overcome this, we employed MVPA to integrate information from across all the sensors and time points (unconstrained decoding), or across all the sensors at each time point (time-series decoding). Unconstrained MVPA provides a sensitive method of assessing effects occurring at any location or combination of locations, and at any time, without increasing type I error (only one statistical test is performed on the pattern of response across all sensors and time points) at the cost of specificity regarding where and when the effect arises (e.g., Grootwagers et al., 2018). Time-series decoding provides more information regarding the timing of discriminant neural patterns, but requires correcting for multiple statistical tests (one per time point).

With our multivariate approach, semantic condition could be decoded from the Neuroscan data in all but two individuals in Experiment 2 (88% detection rate). In our pursuit of an individualised neural marker of language comprehension, this is encouraging. It also confirms the intuition that considering the signal recorded by all electrodes can be more powerful than restricting analysis to one or a few. Even though the N400 effect is well established as having a centro-parietal topography at the group level, our data suggest that there is variation in topology at the individual subject level, and we anticipate that the variation may be even greater amongst minimally-verbal autistic children whose neural patterns are known to be heterogeneous (Salmond et al., 2007). Moreover, we do not want to be too restrictive *a priori* because ultimately, any statistically significant decodable difference between the brain responses to identical auditory tokens presented in different semantic contexts will be meaningful, independent of *where* (and to a certain extent *when*) this effect occurs.

Following this logic, it is also possible that the higher MVPA detection rate obtained with our sentence paradigm compared to the word pairs paradigm reflects neural processes in response to the task (e.g., participants counting – or recognising and attempting to count - the incongruent sentences). In this case, we would not be detecting the brain responses to semantic condition *per se* but rather the brain responses to the task. For our purpose of detecting receptive language however, this would still be meaningful. This is because differential neural processing in one condition indicates that the context of the target word must have been sufficiently processed to influence the way in which the identical auditory token is responded to.

Nonetheless, we acknowledge two limitations of the current approach for future clinical testing. First, MVPA does not specify the nature of the differential processing (where and when it occurred, for the unconstrained approach, or the direction of the effect). Particularly for participants where only a few time points show above-chance decoding accuracy, even though they are corrected for multiple comparisons, we may not feel confident in using this result alone for clinical assessment. As such, it is probably preferable to combine the sensitivity of the MVPA approach with specific and illustrative univariate analyses to increase confidence in the presence of neural signals that reflect language comprehension. Second, for our sentence paradigm where participants had to count the number of incorrect sentences, if we are only decoding the cognitive process of counting, we may miss semantic processing in individuals that lack the cognitive resources to count despite otherwise understanding the sentences.

A critical consideration that follows from this is whether our task will be feasible for the populations we would like to use it with. Although in some regards a completely passive task would be preferable, here we considered a trade-off between the likely sensitivity of the task and how demanding that task is. We hypothesised that at least some minimally-verbal autistic individuals with good receptive language may be able to follow instructions covertly, even if they cannot do so overtly. We do acknowledge, however, that this approach risks missing effects in children who are unable to follow these instructions, perhaps due to poor working memory, attention, or motivation, despite preserved linguistic processing. Finally, we assessed the sensitivity of low-cost wireless EEG technology, Emotiv EPOC+ for detecting N400 effects. For univariate analyses, EPOC+ was able to record N400 effects at the group level and showed a similar detection rate to the research system at the individual subject level. Moreover, when we formally compared the data between systems, we found that the waveforms recorded by EPOC+ were fairly similar in shape and amplitude to those recorded by the research system, with ICC correlation between systems significant, and of “fair” magnitude. Our result adds to the previous literature demonstrating the usability of the EPOC+ to record ERPs (Badcock et al., 2015, 2013; de Lissa et al., 2015a; Duvinage et al., 2013) and shows that late ERPs such as the N400 can also be recorded with a low-cost, portable system. Nonetheless the effects recorded with EPOC+ tended to be numerically lower amplitude than for Neuroscan, and detection rate did not improve with MVPA meaning that the best detection rate for Neuroscan (88%, MVPA, Exp 2) was far above the best with EPOC+ (50%, univariate, Exp 2). Together, our results suggest that Neuroscan would be a preferable option to record quality EEG data, but portable systems such as EPOC+ may prove a valuable alternative in cases where standard EEG setups are unfeasible, such as for certain clinical populations.

A few limitations of the EPOC+ system could be mitigated in future research. First, due to a limitation of the software (Testbench software), the impedance of the EPOC+ electrodes was not assessed precisely and we were only able to ensure that it remained lower than 20 kΩ for each electrode. It is thus likely that impedances were higher for EPOC+ than for Neuroscan (for which impedances were adjusted to < 5 kΩ), although the differences between the systems (e.g., active versus passive electrodes respectively) makes differences in impedance difficult to interpret. Use of a more precise measure of impedance to ensure the best possible connection in every participant would be beneficial. Second, the N400 effect was centrally distributed on the scalp, at least at the group level, in accordance with previous literature (Friedrich & Friederici, 2004; Kutas & Federmeier, 2011), and the EPOC+ headset does not have any central sensors. In future research, researchers should consider wiring an additional electrode in a centro-parietal location where the effects were the largest. It is also important to consider the trade-off between the sensitivity we want to achieve (in which case Neuroscan may be more suitable), and the level of portability and accessibility (in which case EPOC+ is more suitable). In particular, when testing children or special populations, the possibility of recording EEG outside of the lab with an easy and fast setup procedure should be considered as a trade-off against EPOC+’s apparently lower sensitivity, especially in the context of multivariate analyses, and may motivate using portable systems primarily in cases where it would be impossible to obtain data on a research-grade system.

Taken together, our results indicate that it may be possible to index lexical-semantic processing in individual children using EEG. However, contrasting the results we obtained from different paradigms, EEG systems, and analysis methods, several trade-offs have to be considered. The best individual-subject detection rate (88%) was yielded from MVPA of research-grade EEG data in response to identical spoken words presented in the context of congruent and incongruent sentences and a sentence counting task (Experiment 2). Our variable individual subject data emphasise the importance of analysing and reporting individual ERP results, in addition to the grand average data, to illustrate the variability in the presence, location, and timing of ERPs.

## 5 Conclusion

In this study, we set out to establish the rate with which we could detect discriminative brain signals in response to lexico-semantic violations in individual children. We devised two paradigms that used identical spoken language tokens presented in congruent or incongruent lexical-semantic contexts. Additionally, with clinical applications in mind, we tested whether we could use a portable and low-cost device, Emotiv EPOC+, to measure these neural signals. Our results suggest that large inter-individual variability in the neural signatures of lexico-semantic processing exist, even in the neurotypical population. Despite this variability, we replicated group-level N400 effects in neurotypical children using both the EPOC+ and the research-grade Neuroscan system. At an individual level, an N400 effect was evoked in about half of neurotypical children using either Neuroscan or EPOC+ systems. MVPA analyses allowed us to reach near-perfect detection rate with Neuroscan EEG in Experiment 2, with only two participants not showing a reliable electrophysiological signature to semantic anomalies in sentences. Despite limitations for clinical application in its current form, these results give us a basis for future research developing a test for receptive language processing in people who are unable to communicate.

## Author contributions

Conceptualisation: AW, ANR, NAB, JB, LN; Stimulus and task development: AW, NAB, ANR, DM, ND, SY, ES, LN, SP; Data acquisition: SP, AW, NAB; Formal analysis: SP, AW, TG; Writing (original draft): SP; Writing (reviewing and editing): all authors; Supervision: AW.

## 6 Acknowledgments

This work was funded by an Australian Research Council (ARC) Centre of Excellence Neural Markers Training Scheme grant to A.W., N.A.B., A.N.R., J.B and L.N. A.W. was supported by an ARC Future Fellowship (FT170100105) and MRC intramural funding SUAG/052/G101400. L.N. was supported by an ARC Future Fellowship (FT120100102).

We thank Polly Barr and Nickolas Williams for help with data acquisition.

## Footnotes

1 Note that in the Emotiv software, TestBench 3.1.21, the electrodes at FC3/4 are labelled F3/4, F3/4 are labelled AF3/4 and FT7/8 are labelled FC5/6; this is because we adjusted the electrode placement to accommodate the concurrent setup. In this paper we refer to the electrodes according to their placement on the scalp when worn concurrent with the Neuroscan EasyCap, not the labels used in the Emotiv software.

