## Supplementary Table 2 for "Towards an individualised neural assessment of receptive language in children"

Table S2: Stimuli used in Experiment 2: Congruent and incongruent sentences

| **Incongruent sentence frame** | **Congruent sentence frame** | **Target** |
| --- | --- | --- |
| At night the old woman locked the | She plays the guitar so she joined the | Band |
| I roasted marshmallows over the | She saved some money in her piggy- | Bank |
| In the quiet cinema the man's phone | When the thief entered the house the dog | Barked |
| The children ran outside to | In spring flowers | Bloom |
| She jumped in the pool and made a big | The student went to the library to read a | Book |
| His father is the coach of the football | She went to the bakery for a loaf of | Bread |
| He washed his hands with water and | For his wife's birthday he baked a | Cake |
| In autumn leaves fall off the | There were candles on the birthday | Cake |
| Mother hung the painting up on the | The farmer spent the morning milking the | Cows |
| She is afraid of the sea because she cannot | The movie was so sad it made me | Cry |
| In spring flowers | When babies are hungry they often | Cry |
| I put the kettle on to make a hot cup of | The dentist says to brush your teeth twice a | Day |
| She washed the dirty dishes in the | I prefer cats rather than | Dogs |
| The clouds are high up in the | I had no key to open the | Door |
| Most cats see very well at | At night the old woman locked the | Door |
| Today the baby spoke his first | She got out of the car and closed the | Door |
| The letter was sent by | For breakfast he ate scrambled | Eggs |
| There were candles on the birthday | She put on her sunglasses to protect her | Eyes |
| She went to the bakery for a loaf of | They raised pigs on their | Farm |
| The hungry baby wanted to drink | I roasted marshmallows over the | Fire |
| I had no key to open the | He went to the lake to catch | Fish |
| She waited for the phone to | The baby bird was ready to learn to | Fly |
| She accidentally tripped and fell down the | It was windy enough to fly a | Kite |
| He took his dog out for a | At dinner he cut his steak with a | Knife |
| She wore a necklace around her | The hungry baby wanted to drink | Milk |
| He looked up at night to see a sky full of | He was tired and in a bad | Mood |
| When driving you should keep your eyes on the | To hang the picture you need a hammer and | Nail |
| She saved some money in her piggy- | I could not remember his | Name |
| The princess may someday become a | She wore a necklace around her | Neck |
| He cannot post the letter without attaching a | The baby birds were in the | Nest |
| The student went to the library to read a | Most cats see very well at | Night |
| I had no umbrella as it began to | The children ran outside to | Play |
| For breakfast he ate scrambled | The letter was sent by | Post |
| At dinner he cut his steak with a | The princess can only marry a | Prince |
| The baby birds were in the | The princess may someday become a | Queen |
| The movie was so sad it made me | I had no umbrella as it began to | Rain |
| When the thief entered the house the dog | In the quiet cinema the man's phone | Rang |
| The baby bird was ready to learn to | She waited for the phone to | Ring |
| The dentist says to brush your teeth twice a | When driving you should keep your eyes on the | Road |
| It was windy enough to fly a | She washed the dirty dishes in the | Sink |
| I prefer cats rather than | The clouds are high up in the | Sky |
| She put on her sunglasses to protect her | He washed his hands with water and | Soap |
| She forgot her watch so she asked for the | She jumped in the pool and made a big | Splash |
| He went to the lake to catch | She accidentally tripped and fell down the | Stairs |
| THE princess can only marry a | He cannot post the letter without attaching a | Stamp |
| She got out of the car and closed the | He looked up at night to see a sky full of | Stars |
| When babies are hungry they often | She is afraid of the sea because she cannot | Swim |
| The genie promised to grant the man one | I put the kettle on to make a hot cup of | Tea |
| They raised pigs on their | His father is the coach of the football | Team |
| She plays the guitar so she joined the | The dentist opened his mouth to check his | Teeth |
| I could not remember his | She forgot her watch so she asked for the | Time |
| The farmer spent the morning milking the | In autumn leaves fall off the | Trees |
| The dentist opened his mouth to check his | He took his dog out for a | Walk |
| He was tired and in a bad | Mother hung the painting up on the | Wall |
| To hang the picture you need a hammer and | The genie promised to grant the man one | Wish |
| For his wife's birthday he baked a | Today the baby spoke his first | Word |
