## Supplementary Table 1 for "Towards an individualised neural assessment of receptive language in children"

Table S1: Stimuli used in Experiment 1: normatively associated word pairs

| **Unrelated prime** | **Related prime** | **Target** |
| --- | --- | --- |
| House | Leg | Arm |
| Suds | Crafts | Art |
| Knob | Front | Back |
| Sap | Spine | Back |
| Meow | Bounce | Ball |
| Pork | Sand | Beach |
| Spoon | Nest | Bird |
| Seat | Chirp | Bird |
| Quack | Row | Boat |
| Arm | Ship | Boat |
| Filth | Text | Book |
| Cat | Scout | Boy |
| Soil | Girl | Boy |
| Oak | Groom | Bride |
| Stone | Comb | Brush |
| South | Meow | Cat |
| Job | Dog | Cat |
| East | Seat | Chair |
| Rake | Wear | Clothes |
| Nest | Cob | Corn |
| Lamp | Moo | Cow |
| Sand | Calf | Cow |
| Front | Mum | Dad |
| King | Soil | Dirt |
| Quiz | Filth | Dirt |
| Bulb | Pet | Dog |
| Ship | Cat | Dog |
| Calf | Hinge | Door |
| Bride | Knob | Door |
| Girl | Quack | Duck |
| Groom | Bait | Fish |
| Boy | Tile | Floor |
| Queen | Spoon | Fork |
| Moo | Back | Front |
| Clock | Boy | Girl |
| Bed | Vine | Grape |
| Text | Bride | Groom |
| Tile | Glove | Hand |
| Chirp | Cap | Hat |
| Cob | House | Home |
| Scout | Honk | Horn |
| Thigh | Queen | King |
| Wear | Rake | Leaves |
| Cap | Arm | Leg |
| Nap | Thigh | Leg |
| Mum | Bulb | Light |
| Pet | Lamp | Light |
| Honk | Dad | Mum |
| Black | South | North |
| Row | Ink | Pen |
| Glove | Pork | Pig |
| Dad | Stone | Rock |
| Gums | Socks | Shoes |
| Ink | Bed | Sleep |
| Vine | Nap | Sleep |
| Crafts | Suds | Soap |
| Comb | North | South |
| Socks | Gums | Teeth |
| Spine | Quiz | Test |
| Leg | Clock | Time |
| Bait | Sap | Tree |
| Hinge | Oak | Tree |
| Dog | East | West |
